# Molecular characterisation of *Streptococcus pyogenes* (StrepA) non-invasive isolates during the 2022-23 UK upsurge

**DOI:** 10.1101/2024.05.02.592214

**Authors:** Jennifer N. Hall, Saikou Y. Bah, Henna Khalid, Alison Brailey, Sarah Coleman, Tracey Kirk, Naveed Hussain, Mark Tovey, Roy R. Chaudhuri, Steve Davies, Lisa Tilley, Thushan de Silva, Claire E. Turner

## Abstract

At the end of 2022 into early 2023 the UK Health Security Agency reported unusually high levels of scarlet fever and invasive disease caused by *Streptococcus pyogenes* (StrepA or group A *Streptococcus*). During this time, we collected and genome sequenced 341 non-invasive throat and skin *S. pyogenes* isolates identified during routine clinical diagnostic testing in Sheffield, a large UK city. We compared the data with that obtained from a similar collection of 165 isolates from 2016-17.

Numbers of throat-associated isolates collected peaked in early December 2022, reflecting the national scarlet fever upsurge, while skin infections peaked later in December. The most common *emm*-types in 2022-23 were *emm*1 (28.7%), *emm*12 (24.9%), and *emm*22 (7.7%) in throat; and *emm*1 (22%), *emm*12 (10%), *emm*76 (18%), and *emm*49 (7%) in skin. Whilst all *emm*1 isolates were the M1_UK_ lineage, comparison with 2016-17 revealed diverse lineages in other *emm*-types, including *emm*12, and emergent lineages within other types including a new acapsular *emm*75 lineage, demonstrating that the upsurge was not completely driven by a single genotype. Analysis of the capsule locus predicted only 51% of throat isolates would produce capsule compared to 78% of skin isolates. 90% of throat isolates were also predicted to have high NADase and Streptolysin O (SLO) expression, based on the promoter sequence, compared to only 56% of skin isolates.

Our study has highlighted the value in analysis of non-invasive isolates to characterise tissue tropisms, as well as changing strain diversity and emerging genomic features which may have implications for spillover into invasive disease and future *S. pyogenes* upsurges.

**Data summary:** All new genome sequence data is available on the NCBI short read archive under the bioproject PRJNA1062601 and individual accession numbers are listed in **Supplementary Table 1 and Table 2**.

**Impact statement:** The human bacterial pathogen *Streptococcus pyogenes*, also known as group A *Streptococcus* or StrepA, caused a dramatic and sudden upsurge in scarlet fever in the UK at the end of 2022 into early 2023. We present molecular characterisation of this upsurge, through genome sequence analysis of throat, skin and other types of non-severe infection isolates collected by the microbiology diagnostic lab at the Northern General Hospital in Sheffield, England. We found that, whilst two strain types were the predominant cause of infections during the upsurge, other types had emerged or changed when compared to a similar collection from 2016-17. We also identified differences between throat-associated isolates and skin-associated isolates and highlighted important bacterial factors that might influence infection types. Isolates from non-severe throat/skin types of infections are rarely saved and therefore our knowledge of them is limited. However, here we demonstrate that study of such isolates may be key to understanding upsurges of more severe infections.

## Introduction

The human pathogen *Streptococcus pyogenes*, also known as group A *Streptococcus* (GAS) or StrepA, is a common cause of throat infections, such as pharyngitis and tonsillitis, as well as mild skin infections such as pyoderma or impetigo. More commonly in children than adults, throat infections can progress to scarlet fever, with a characteristic sandpaper-like rash and ‘strawberry tongue’. On rare occasions, *S. pyogenes* can also cause severe and potentially lethal invasive diseases, such as pneumonia, empyema, bacteraemia and necrotising fasciitis (‘flesh-eating’ disease). In England, scarlet fever and invasive *S. pyogenes/*group A *Streptococcus* (iGAS) disease cases are notifiable to the UK Health Security Agency (UKHSA).

In September 2022, UKHSA reported an unusually high level of scarlet fever notifications with ∼3.7 fold more than in the same period for the previous five seasons [1]. Notifications continued to rapidly increase across England and Wales, with 8,688 cases in weeks 37-48 (mid-Sept to end-Nov), compared to 333-2,536 in the previous five seasons [2]. Alongside the increasing scarlet fever cases, there were also high numbers of iGAS notifications with 772 in weeks 37-48. Concerningly, during these weeks there were more cases of iGAS in children under 15 (26.1%), compared to previous seasons (6.4-13.3%) and 14 deaths in this age group at this time [2]. Scarlet fever notifications peaked in week 49 with 10,069 cases, and iGAS notifications peaked in week 52 with 213 cases [3]. Typing of isolates by the sequence of the *emm* gene, which encodes for the hypervariable M protein, identified *emm*1 as the most common cause of iGAS in those older than 15 (31%) but an even higher proportion of cases in those younger than 15 (57%). *emm*12 and *emm*4 were the second (23%) and third (7%) most common causes of iGAS in children, at least in the early part of the upsurge [2].

Scarlet fever demonstrates seasonality with cases typically increasing in late winter and peaking in early spring. An unexpected increase in scarlet fever cases in England was first seen in 2013-14, peaking in early April and totalling over 13,000 notifications, compared to fewer than 3000 in previous seasons [4,5]. Notifications remained seasonally elevated, rising each year to their highest in 2017-18 [6], until the COVID-19 pandemic when cases fell dramatically. No corresponding rise was observed for iGAS notifications until the 2015-16 season. The link between scarlet fever cases and iGAS cases is not well understood but it appeared that the increase in both scarlet fever and invasive disease notifications in early 2016 was due to the emergence of a new variant of *emm*1 termed M1_UK_ [7,8], which, prior to 2022-23, also led to the biggest upsurge in cases in 2017-18 [6]. The M1_UK_ variant had been steadily increasing in prevalence in England since 2010 and represented 91.5% of invasive *emm*1 by 2020 [8]. M1_UK_ is characterised by 27 SNPs in comparison to a globally circulating *emm*1 population (M1_global_), and increased expression of the superantigen *speA* [7,9,10]. The M1_UK_ lineage has been detected in European countries, North America, Australia and New Zealand, often associated with increases in disease [7,9,11–17].

Whilst iGAS undergoes routine surveillance in several high-income countries, with the inclusion of scarlet fever and outbreak situations in some, these types of infections only represent a small proportion of streptococcal cases. Our knowledge of circulating non-invasive disease (from non-sterile sites) isolates is severely lacking, yet these isolates may act as an early indicator for increasing prevalence of new genotypes or lineages that could lead to more severe infections. This lack of knowledge also means we have a limited understanding of the connection between certain *emm*-types and preferences for causing throat infections or skin infections. As non-invasive isolates are not routinely collected, data acquisition relies on local collections.

Sheffield is a large northern city in England with a population of around 600,000. During the 2022-23 upsurge the region of Yorkshire and the Humber, which includes Sheffield, reported the highest rates of invasive *S. pyogenes* in England, at 8.7 per 100,000 population [3]. Scarlet fever notifications were also high in the region, at 132.0 per 100,000 population, although this was similar to the rate seen in the North West region and lower than the East Midlands region [3]. We began in November 2022 to routinely save all non-invasive isolates identified by the Department of Laboratory Medicine at the Northern General Hospital, Sheffield, that performs microbiological services for community care as well as surrounding hospitals. We had also previously performed a similar collection in 2016-17 which we used as a comparative population. We undertook whole genome sequence analysis of both collections and characterised strain diversity and pathogenicity factors. As expected, the *emm*1 M1_UK_ lineage dominated during the 2022-23 upsurge followed by *emm*12, but there were some unexpected increases in other *emm*-types as well as differences between throat-associated isolates and skin-associated isolates.

## Methods

### Isolate collection

A total of 384 non-invasive isolates (from non-sterile sites), presumptively identified from culture as *S. pyogenes*, were collected from the Department of Laboratory Medicine, Northern General Hospital, Sheffield, United Kingdom, between November 2022 (week 45) and February 2023 (week 6). The Department of Laboratory Medicine performs microbiology diagnostics for NHS Trusts as well as primary care and community services, acting as the single regional diagnostic microbiology laboratory for a population of around 600,000 adults and children in Sheffield. Anonymised clinical data were collected for each isolate from the sample request information and electronic clinical patient records: swab source (throat/skin/ear/eye/nose), sampling date, age, sex, and infection type. Cases were considered to be associated with scarlet fever where scarlet fever was queried by the clinician or a rash consistent with a scarlet fever diagnosis was described in the clinical details accompanying the request. All other throat samples, and other sample types, were deemed non-scarlet fever samples. Samples from ear, eye and nose isolates were excluded from further downstream analyses to focus the comparison on throat and skin isolates.

For comparison, a further 229 archived non-invasive *S. pyogenes* isolates collected in a similar manner between October 2016 and January 2017 were also included in this study, with data collected as above. The 2016-17 season also showed a seasonal upsurge in disease compared to previous years, however with substantially fewer scarlet fever and invasive *S. pyogenes* cases reported compared to 2022-23. The age distribution of cases in 2016-17, of both scarlet fever and invasive *S. pyogenes* disease, was consistent with previous years and this was therefore considered a suitably representative comparative sample.

### Whole-genome sequencing (WGS)

Genomic DNA was extracted from all isolates using a previously described method [18]. Isolates collected in 2022-23 underwent whole genome sequencing (WGS) at Earlham Institute by the Genomics Pipeline group, using the LITE protocol [19] for library preparation and sequenced on a NovaSeq X plus generating 150bp PE reads. Isolates collected in 2016 underwent sequencing provided by MicrobesNG (https://microbesng.com) using the Nextera XT library prep kit (Illumina) and the Illumina HiSeq 2500, generating 250-bp paired-end reads.

### WGS analysis

Raw sequence reads were trimmed using Trimmomatic (v0.39) with the settings LEADING:3 TRAILING:3 SLIDING WINDOW:4:15 MINLEN:36 [20]. For the 2022-23 isolates, the average estimated read coverage was ∼524x with some greater than 1000x. Reads were therefore randomly subsampled, using seqtk, to 1.2 million reads per isolate, to provide ∼180x coverage.

Trimmed and subsampled reads were then used to perform *de novo* assembly using SPAdes (v.3.13.1) with k-mer sizes of 21, 33, 55 and 77 [21]. Assembly statistics were generated for each isolate using Quast [22] (**Supplementary Tables 1 and 2**) and any draft assemblies with more than 500 contigs or a total genome size greater than 2.2 Mb were excluded from downstream analysis, as were any that were determined not to be *S. pyogenes*. Multi-locus sequence types (MLST) and *emm*-types were determined from the *de novo* assemblies using mlst (https://github.com/tseemann/mlst) with the pubmlst database [23], and the emm_typer.pl script (github.com/BenJamesMetcalf/GAS_Scripts_Reference), respectively. New MLSTs were submitted to pubmlst and new *emm*-types and sub-types to the CDC *emm*-type database (https://cdc.gov/streplab) for assignment.

*De novo* assemblies were annotated using Prokka (v.1.14.6) [24]. Snippy (https://github.com/tseemann/snippy) was used to determine single nucleotide polymorphism (SNP) distances between sequence reads and a reference genome. RAxML v8.2.12 [25] was used to generate maximum likelihood phylogenetic trees based on the core gene alignment with a general time-reversible (GTR) substitution model and 100 bootstraps. In some instances, regions of predicted recombination were identified and removed using Gubbins [26] prior to tree construction. Phylogenetic trees were annotated using iTOL (version 6) [27].

Further comparison was made with previously published WGS data from Sheffield, consisting of 142 non-invasive skin and soft tissue isolates collected in 2019 [28], and national and international data from other published collections.

### Variable factor typing

The presence of superantigen genes *speA, speC, speG, speH, speI, speJ, speK, speL, speM, speK/M, speQ, speR, ssa*, and *smeZ* and DNase genes *sda1, sda2, sdn, spd1, spd3*, and *spd4* were determined by BLAST (100% coverage, 80% identity) with representative gene sequences against assemblies, and a manual check where needed. For other genes (*covR/S*, *rocA*, *hasABC* and the *nga-ifs-slo* locus with promoter region), sequences were extracted from the assembled genomes and compared to the reference genome, H293 (Genbank accession NZ_HG316453.1), as previously described [28]. Antimicrobial resistance (AMR) gene carriage was determined with ABRicate (https://github.com/tseemann/abricate) using the NCBI database [29].

### Ethical approval

The study protocol was approved by a Research Ethics Committee (reference 24/NE/0033; Integrated Research Application System project ID 334500).

## Results

### Collected isolates

During the unusual *S. pyogenes* infection upsurge period reported by UKHSA in England 2022-23, we collected a total of 384 non-invasive isolates from the Department of Laboratory Medicine, Sheffield. This was compared to 229 isolates collected in 2016-17, representing a time of more typical seasonal upsurge. The proportion of throat isolates were similar for both time periods (61.7% and 65%) but during 2022-23 overall isolate numbers were much higher (**Table 1**). Significantly more isolates were also associated with scarlet fever in 2022-23 than in 2016-17 (24.5% vs 5.4%, χ^2^(1) = 23.55, p<0.0001). For both time periods, throat isolates were primarily from children under 9, but in 2022-23 significantly more were from children aged 5-9 compared to 2016-17 (33.3% vs 15.7%, χ^2^(1) = 22.00, p<0.0003). There were fewer skin than throat isolates for both time periods, and skin isolates were predominantly from those aged one year and under, and those aged 45 and older. There was evidence in both time periods of a bimodal age distribution with the highest number of isolates from children under 9 years of age, and a second peak in individuals aged 30-39 years (**Supplementary Figure 1**). *S. pyogenes* was more frequently isolated from throat swabs in female patients (57.6% (136/236) in 2022-23 and 63.8% (95/149) in 2016-17) and from skin swabs in male patients (68.8% (77/112) in 2022-23 and 63.2% (43/68) in 2016-17) (**Supplementary Figure 1**).

**Table 1:** Clinical characteristics of isolates collected in 2022-23 and 2016-17.

|  | 2022-23 (%) | 2016-17 (%) |
| --- | --- | --- |
| **Clinical Presentation** |  |  |
| Throat | 237 (61.7) | 149 (65.0) |
| - Scarlet fever | 58 (24.5) | 8 (5.4) |
| - No Scarlet fever | 179 (75.5) | 141 (94.6) |
| Skin | 111 (28.9) | 68 (29.7) |
| Other | 36 (9.4) | 12 (5.2) |
| - Ear | 28 (7.3) | 10 (4.4) |
| - Eye | 7 (1.8) | 0 (0) |
| - Nose | 1 (0.3) | 2 (0.9) |
| **Sex** |  |  |
| Female | 190 (49.5) | 125 (54.6) |
| Male | 194 (50.5) | 104 (45.4) |
| **Age<sup>#</sup>** |  |  |
| 1 year and under | 26 (6.8) | 10 (4.4) |
| 2-4 years | 64 (16.7) | 38 (16.6) |
| 5-9 years | 128 (33.3) | 36 (15.7) |
| 10-14 years | 33 (8.6) | 20 (8.7) |
| 15-44 years | 97 (25.3) | 103 (45.0) |
| 45-64 years | 22 (5.7) | 13 (5.7) |
| 65-74 years | 9 (2.3) | 5 (2.2) |
| 75 years and over | 5 (1.3) | 4 (1.7) |
| **Total** | **384 (100)** | **229 (100)** |

<sup>#</sup>Isolates grouped in accordance with UKHSA age categories.

### Emm-type distribution

After quality control filtering, a total of 341 whole genome sequences from the *S. pyogenes* isolates collected in 2022-23 were included in study analyses: 209 (61.3%) from throat swabs, 54 (25.8%) of which were associated with scarlet fever; 100 (29.3%) from skin swabs; and 32 (9.4%) from other sites. From the 2016-17 collection, 165 isolates passed quality control filtering: 127 (77.0%) from throat swabs, and 38 (23.0%) from skin swabs. We purposely did not sequence isolates from ‘other’ sites in this earlier collection.

The *emm*-type for each isolate was extracted from the WGS data. The frequency of isolates and the distribution of *emm*-types varied over time in the 2022-23 collection, with the total number peaking in week 49, in keeping with UKHSA data [3]. We observed a fall in the quantity of throat swabs following the issue of interim clinical guidance by NHS England on 9th December 2022 (week 49) which temporarily altered the empirical treatment threshold for presumed streptococcal throat infections. Across all 2022-23 isolates, a total of 33 different *emm*-types were identified, with *emm*1 being the most common at 95/341 (27.9%) followed by *emm*12, at 64/341 (18.8%) (**Figure 1A**). This high level of *emm*1 and *emm*12 cases was reflected in an increase in the number of throat samples in late November - early December 2022 (**Figure 2**). Within our comparative collection from 2016-17, 28 different *emm*-types were identified overall, most frequently *emm*89 (30/165, 18.2%).

**Figure 1:**
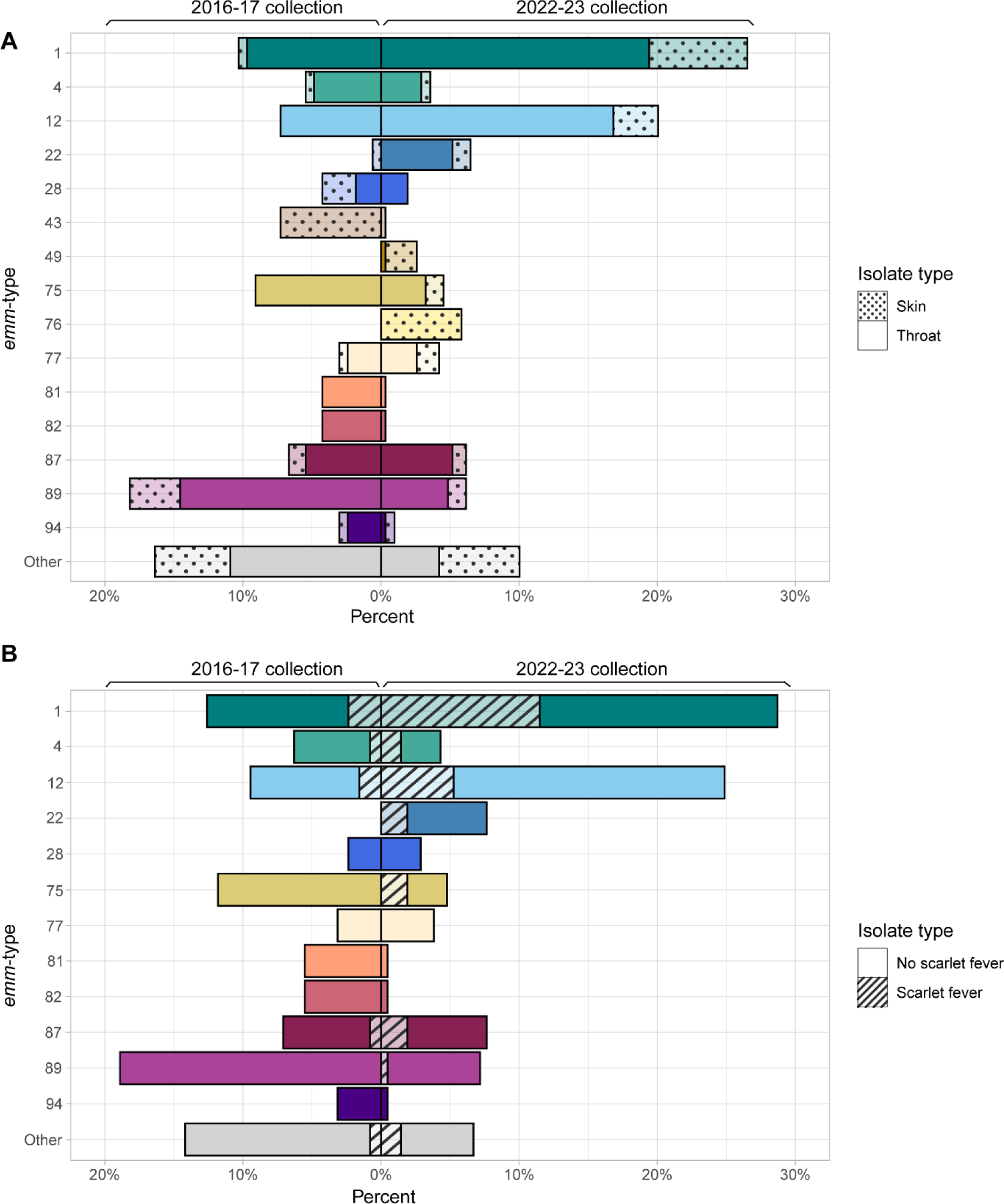
(A) Distribution of *emm*-types by collection year and clinical isolate type. (B) Distribution of *emm*-types within throat isolates by clinical presentation. Data presented as a percentage of (A) the total number of isolates and (B) the total number of throat isolates in each collection period.

**Figure 2:**
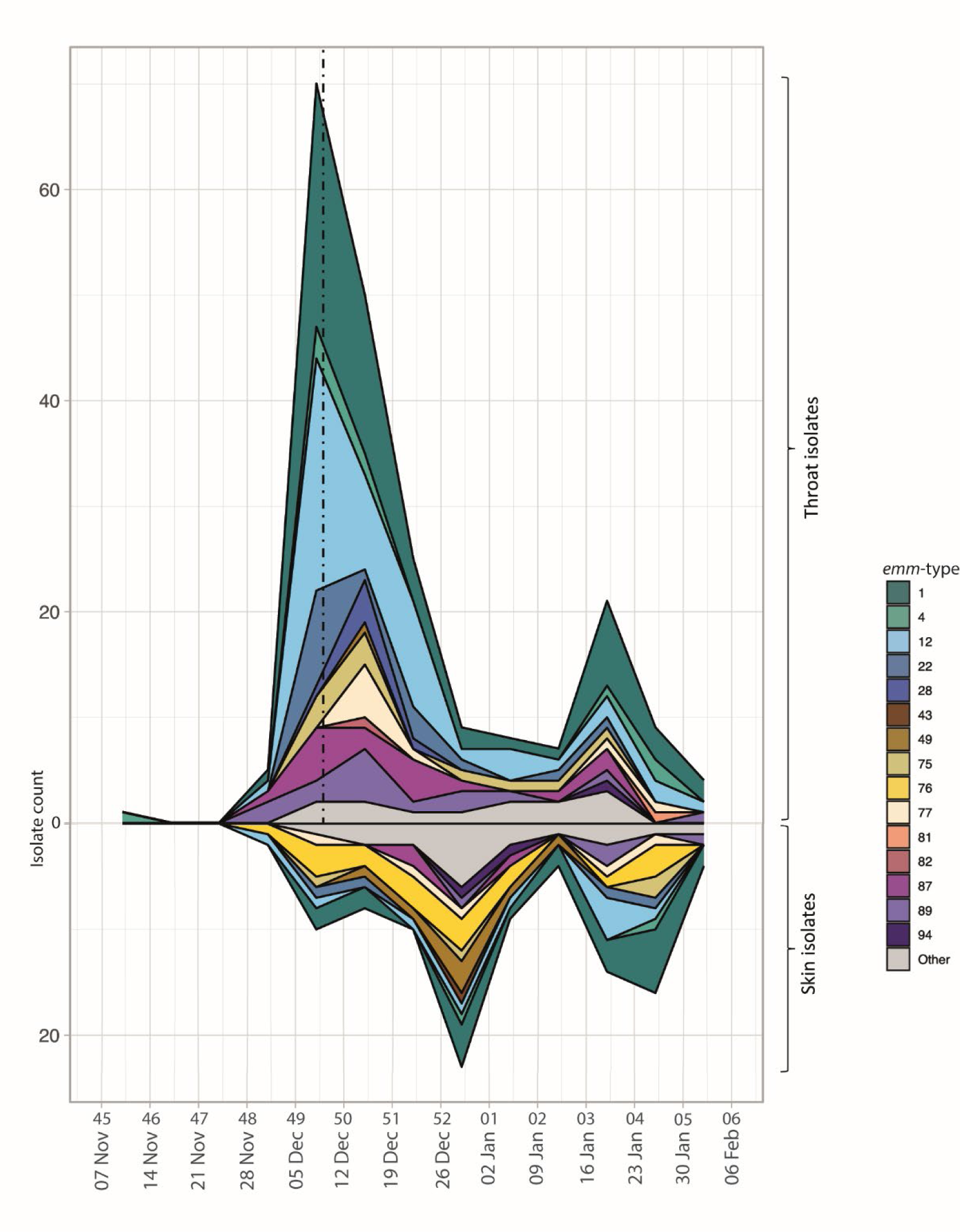
Distribution of *emm*-types across time for throat and skin isolates collected in 2022-23. Week numbers and dates are presented on the x axis. The dashed line represents the introduction of the NHS England group A *Streptococcus* interim clinical guidance summary for case management, on 9th December 2022.

Within the 2022-23 throat isolates, 22 different *emm*-types were identified, but just six *emm*-types made up 80.9% of isolates: *emm*1 (28.7%, 60/209), *emm*12 (24.9%, 52/209), *emm*22 (7.7%, 16/209), *emm*87 (7.7%, 16/209), *emm*89 (7.2%, 15/209) and *emm*75 (4.8%, 10/209). *Emm*1 and *emm*12 made up a significantly higher proportion of throat isolates in 2022-23 compared to the comparative collection from 2016-17, where 12.6% (16/127) isolates were *emm*1 (χ^2^(1) = 10.811, p = 0.001) and 9.5% (12/127) were *emm*12 (*χ^2^*(1) = 11.22, p = 0.0008). In 2022-23 throat isolates, *emm*89 made up a significantly lower proportion than the 2016-17 collection, in which it was the most common *emm*-type at 18.9% (24/127) isolates (*χ^2^*(1) = 9.4656, p = 0.002). There was also a lower proportion of *emm*75 within the 2022-23 throat isolates compared to 2016-17, where *emm*75 made up 11.8% (15/127). *Emm*87 was a similar proportion in 2016-17 (7.1%, 9/127), however no throat isolates were *emm*22 in 2016-17. Throat isolates associated with scarlet fever in 2022-23 were predominantly *emm*1 (44.4%, 24/54) and *emm*12 (20.4%, 11/54) (**Figure 1B**). In 2016-17, scarlet fever was associated with 8/127 throat isolates of five *emm*-types: *emm*1 (3/8), *emm*12 (2/8), *emm*4 (1/8), *emm*6 (1/8), and *emm*87 (1/8).

The 2022 upsurge in throat disease was followed by a surge in skin disease in late December 2022, with a second peak in late January 2023 (**Figure 2**). This was driven by a range of different *emm*-types, with 23 different types overall. Again, frequently *emm*1 at 22/100 (22%) isolates and *emm*12 at 10/100 (10%) isolates, but also *emm*76 at 18/100 (18%) and *emm*49 at 7/100 (7%), both of which were rarely found in the throat. Furthermore, differences in *emm*-type distribution were seen when compared to our comparative collection from 2016-17, in which the dominant skin isolate *emm*-types were *emm*43 (31.6%, 12/38), followed by *emm*89 (15.8%, 6/38) and *emm*28 (10.5%, 4/38) (**Figure 1A**). A total of 17 different *emm*-types were identified within the skin isolates in the 2016-17 collection, with only a single skin isolate identified as *emm*1, and none as *emm*12 or *emm*76.

An *emm*-pattern was assigned to each isolate using previous *emm*-type assigned patterns based on the genes surrounding the *emm*-gene [30]. Within the 2022-23 collection, there was a similar proportion of ‘throat-associated’ *emm*-pattern A-C (48.1%, 164/341) and ‘generalist’ *emm*-pattern E (48.4% 165/341), with just 3.5% (12/341) being ‘skin-associated’ pattern D (**Figure 3**). All pattern D isolates were of skin origin, except a single throat isolate. Within the 2016-17 collection, ‘generalist’ *emm*-pattern E were more common at 62.4% (103/165); 23% (38/165) were pattern A-C. With 14.5% (24/165) of isolates pattern D, the 2016-17 collection had a greater proportion of pattern D than the 2022-23 collection overall, with the most striking difference noted in the skin isolates, where pattern D made up 44.7% (17/38) of the 2016-17 skin isolates but just 11% (11/100) of the 2022-23 skin isolates. Just one skin isolate collected in 2016-17 was of *emm*-pattern A-C. Despite these differences and consistent with their previously defined tissue-tropism, in both time periods *emm*-pattern A-C were significantly more associated with throat isolates than skin (2016-17 *χ*^2^(1) = 11.59, p=0.0007; 2022-23 *χ^2^*(1) = 13.15, p=0.003), while *emm*-pattern D were significantly more associated with skin isolates than throat (2016-17 *χ^2^*(1) = 36.20, p<0.0001; 2022-23 *χ^2^*(1) = 20.06, p<0.0001), and pattern E isolates were not significantly associated with one or the other.

**Figure 3:**
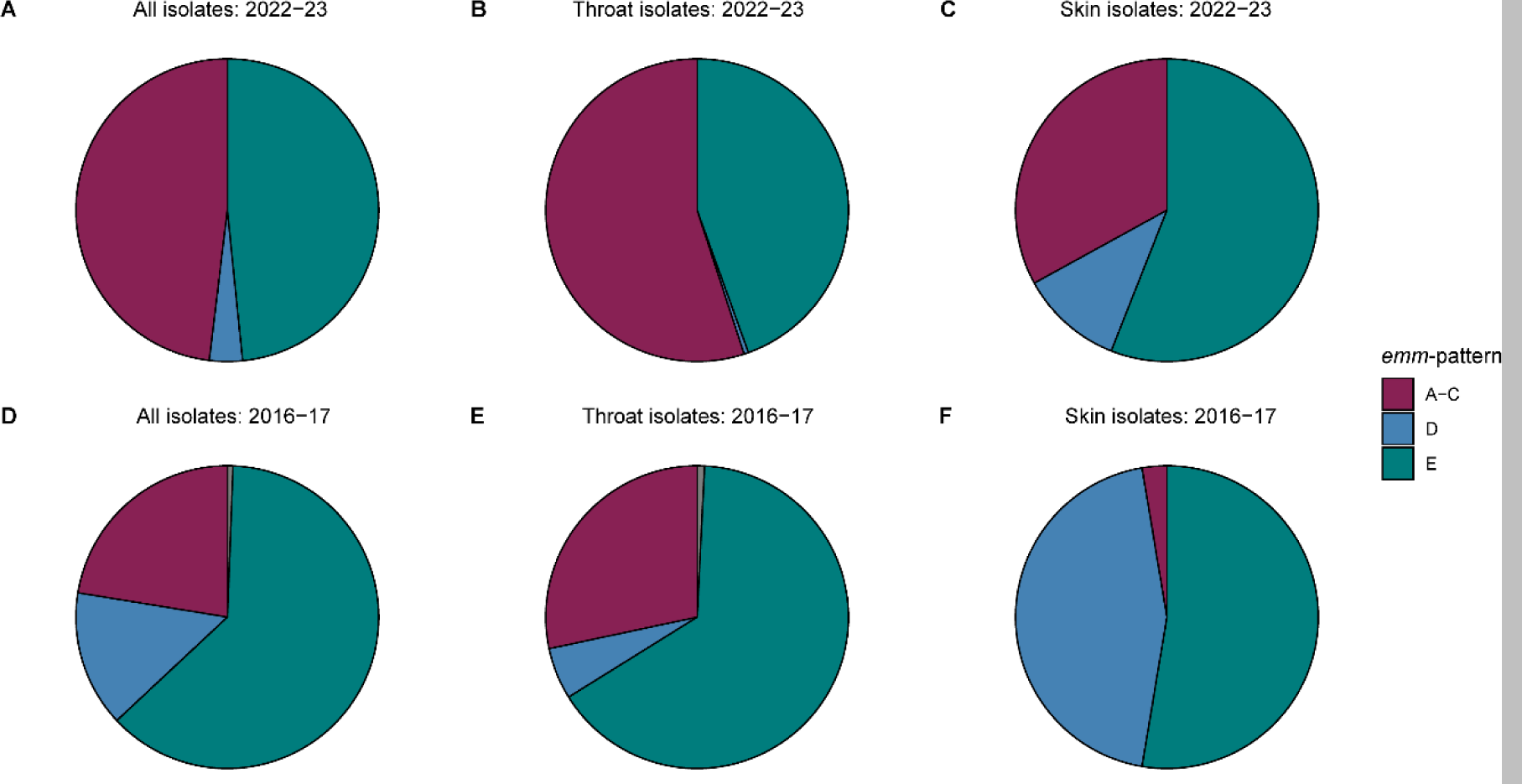
Distribution of *emm*-patterns across 2022-23 and 2016-17 collections. (**A**) all 2022-23 isolates; (**B**) 2022-23 throat isolates; (**C**) 2022-23 skin isolates; (**D**) all 2016-17 isolates; (**E**) 2016-17 throat isolates; (**F**) 2016-17 skin isolates. Pie charts represent the percentage of isolates associated with each pattern.

Each isolate was also assigned an *emm*-cluster (**Supplementary Figure 2**). Within the 2022-23 collection, the most common was the *emm*1 cluster type A-C3 (27.9%, 95/341), and the *emm*12 cluster type A-C4 (18.8%, 64/341). E4 was also common (19.6%, 67/341), representing *emm*22*, emm*77*, emm*89 *and emm*102. By comparison, *emm*-cluster E4 made up the largest proportion of isolates within the 2016-17 collection (28.5%, 47/165), reflecting the high number of *emm*89 isolates from both throat and skin samples. The 2016-17 collection also had a number of samples in cluster D4 (9.7%, 15/165), all of which were skin isolates and mostly *emm*43 (80%, 12/15).

### Antimicrobial resistance genes

The presence of antimicrobial resistance genes in the genomes of our 2022-23 throat and skin isolates was relatively low overall (**Supplementary Table 1**), with only 29.4% (91/309) of isolates carrying at least one gene. By far the most common was *tetM* at 24.3% (75/309) and 28.5% of isolates carried at least one *tet* gene (*tetL, tetM, tetO* or *tetT*). The second most common resistance gene was *ermA* (4.9%). No resistance genes were identified in *emm*1 and only three *emm*12 carried resistance genes (*mefA* and *msrD* encoding macrolide resistance). Fewer 2016-17 throat and skin isolates carried at least one resistance gene (18.2%, 30/165) (**Supplementary Table 2**) but all of these included a *tet* gene (*tetM* or *tetO*), with *tetM* being the most common at 15.8% (26/165).

### Hyaluronic acid capsule synthesis and the nga-ifs-slo toxin loci

We previously identified an increasing number of *emm*-types that had undergone recent genetic changes leading to the inability to produce the hyaluronic acid capsule, through loss or nonsense mutation of capsule synthesis genes *hasABC* [31]. We therefore examined the *hasABC* locus in the genomes of our Sheffield 2022-23 throat and skin isolates and found overall, 40% of isolates were predicted to be unable to synthesise the hyaluronic acid capsule due to mutations in or absence of *hasA, hasB* and *hasC* genes. While sporadic nonsense mutations occurred in *hasA* or *hasB* in some *emm*-types, such as *emm*1 (4.9%) and *emm*12 (14.5%), 100% of all isolates belonging to 11 different *emm*-types were predicted to be acapsular (**Supplementary Table 1**). This included *emm*22 and *emm*89, for which, as expected, the entire *hasABC* locus was absent, and *emm*28, *emm*77 and *emm*87 which have previously described nonsense mutations in *hasA* [31,32]. Typically, *emm*4 also lacked the *hasABC* locus although one isolate was found to carry it. Nonsense *hasA* mutations were also found in the majority of *emm*75 (78.5%) and *emm*11 (66.7%).

Within the 2022-23 throat isolates, only 51% were predicted to be able to produce capsule compared to 78% of skin isolates. Loss of capsule was predominantly associated with *emm*-pattern E isolates, for which only 27% were predicted to be encapsulated compared to 91% and 92% of pattern A-C and D, respectively. This was similar in the 2016-17 isolates, where only 30% of pattern E isolates were predicted to be able to make capsule (**Supplementary Table 2**).

We have previously identified convergent evolution with acapsular isolates also having undergone homologous recombination, resulting in increased expression of the toxins NADase and Streptolysin O (SLO) [31,33,34]. High or low expression of these toxins can be linked to three residues in the promoter region of the *nga* (encoding for NADase), *ifs* (encoding the inhibitor of NADase) and *slo* locus [34]. Within the 2022-23 throat isolates, 90% were predicted to have high-toxin expression (as defined previously [31]), compared to 56% of skin isolates (**Supplementary Table 1**). Only 8% of pattern D isolates were predicted to have high-toxin expression, compared to 99% of A-C and 66% of E. Additionally only 58% of pattern D isolates would express active NADase, based on a glycine residue at codon 330 rather than an aspartate [35], compared to 99% and 100% of pattern A-C and E isolates, respectively. Although *emm*1 and *emm*12 isolates were predominantly encapsulated with a high-toxin expression genotype, an acapsular with a high-toxin genotype was found in 41% of throat isolates compared to a significantly lower 22% of skin isolates (χ^2^(1) = 66.315, p < 0.0001). Only 2% of throat isolates were predicted to be encapsulated with low-toxin expression compared to 44% of skin isolates. This was similar in 2016-17 with 8% of throat isolates predicted to be encapsulated with low-toxin expression and 50% of skin isolates (**Supplementary Table 2**).

### Superantigens

*S. pyogenes* has the potential to carry at least one of 13 different superantigen genes: *speG*, *speJ*, *speQ*, *speR*, and *smeZ* are chromosomal, whilst *speA*, *speC*, *speH*, *speI*, *speK*, *speL*, *speM*, and *ssa* are prophage-associated. Within 2022-23 throat and skin isolates, 96.1% (297/309) carried *smeZ* and 92.2% (285/309) carried *speG* (**Supplementary Figure 3A**). The other chromosomal superantigens were seen less frequently, with *speJ* in 42.4% (131/309) and the co-transcribed *speQ* and *speR* in 12.6% (39/309). Of the prophage-associated superantigens, *speC* was the most common at 52.4% (162/309), followed by *speA* at 32% (99/309). One *emm*49 skin isolate carried a *speK/speM* fusion gene which we previously identified in an *emm*65 [28]. Of the 99 isolates that carried *speA*, 80.8% (80/99) were *emm*1.

In comparison, a similar proportion of isolates from the 2016-17 collection carried the most common superantigen genes, *smeZ* and *speG*, at 94.5% (156/165) and 90.9% (150/165) respectively (**Supplementary Figure 3B**). Prophage-associated *speC* was found in 64.8% (107/165) of isolates in 2016-17, slightly higher than in 2022-23. However, the number of isolates carrying *speJ* and *speA* was lower in 2016-17, where *speJ* was found in 22.4% (37/165) and *speA* in 16.4% (27/165); this difference reflects the 97.6% (80/82) of *emm*1 isolates carrying this combination of superantigens within the 2022-23 collection. In 2022-23, 22% (68/309) carried *speI* compared to 16.4% (27/165) in 2016-17, reflecting the upsurge in *emm*12 in 2022-23, all of which carried *speI*. A lower proportion had *speK* in 2022-23 (14.6%, 45/309) than 2016-17 (26.1%, 43/165), reflecting the single *emm*43 isolate in 2022-23 compared to 12 *emm*43 skin isolates in 2016-17. One *emm*25 skin isolate in 2016-17 carried a *speK/speM* fusion gene.

### DNases

All *S. pyogenes* strains carry at least one DNase gene, out of a possible two chromosomal and six prophage-encoded genes [36]. Within the 2022-23 throat and skin isolates, the most common prophage-associated DNase genes identified were *spd3* in 60.5% (187/309), *spd1* (carried with *speC*) in 52.4% (162/309), and *sda2* in 43.0% (133/309) (**Supplementary Figure 4A**). Most DNase combinations were seen in a range of *emm*-types, although the most common combinations were specific to *emm*1 (*sda2* with *spd3*, 23.6%, 73/309) and *emm*12 (*sda2* with *spd1,* 13.3%, 41/309). By comparison, in the 2016-17 collection *sda2* was seen in fewer isolates, at 21.2% (35/165), and *spd1* in slightly more, at 64.2% (106/165), but *spd3* in similar numbers, at 56.4% (93/165) (**Supplementary Figure 4B**). In both collections, *sdn* was less common and *spd4* was quite rare, with *spd4* only being present with *spd3* in a single isolate in 2022-23 and two isolates in 2016-17 (two *emm*5 and one *emm*87); all other isolates carrying *spd3* or *spd4* had just one of these two DNase genes.

### CovR, CovS and RocA regulator proteins

The two-component regulator CovR/S and the regulator of Cov, RocA, negatively regulate key *S. pyogenes* virulence factors including capsule and toxins. For both collections, nonsense or frameshift mutations leading to premature stop codons that would alleviate virulence factor repression were rare in CovS and RocA and absent in CovR. Only one 2022-23 isolate had a premature stop codon in CovS (an *emm*1), and two 2016-17 isolates (one *emm*1 and one *emm*89). A total of 2.6% (8/309) 2022-23 throat or skin isolates carried premature stop codons that would truncate RocA. Consistent with previous findings, all *emm*3 isolates from both collections would express a truncated RocA after 416 aa. No other *emm*-types had nonsense mutations in *rocA* in 2016-17. Other non-synonymous variations were found in *covS* and *rocA* (**Supplementary Tables 1 and 2**) but the impact of these is difficult to determine and many were associated with *emm*-type.

Although nonsense mutations in *covR* were absent, other non-synonymous variations in *covR* were found in 45/309 (14.6%) of 2022-23 skin and throat isolates. However, the majority were related to the *emm*-type. All *emm*22 isolates from 2022-23 (n=20) and 2016-17 (n=1) had the same variation in CovR (V128A) and also had a variation in RocA (V333A). Nearly all *emm*77 isolates from both 2022-23 (12/13) and 2016-17 (4/5) had the same M170I variation in CovR. 14.5% of 2022-23 *emm*12 isolates (9/62) had an A105G variation in CovR. Four other isolates of different *emm*-types had other CovR variations.

### Emm1

A total of 95 (27.9%) of the 341 genomes collected in 2022-23 were *emm*1. Sixty (63.2%) of these were from throat samples, of which 24 were associated with scarlet fever. Twenty-two (23.2%) were skin isolates, and the remaining 13 from other sites including 12 from ear swabs and one from an eye swab. All of the *emm*1 isolates clustered with the M1_UK_ lineage and all were confirmed to carry the 27 SNPs that define this lineage (**Figure 4**). This was similar to our comparative collection from 2016-17, in which there was only a single M1_global_ isolate out of 17 *emm*1 isolates, with the remaining 16 being M1_UK_. This is in keeping with the emergence of M1_UK_ as the dominant *emm*1 strain globally. All Sheffield strains were ST28, except a single Sheffield 2022-23 *emm*1 skin isolate which was ST1357, differing from ST28 by a single polymorphism in the *gtr* locus. Sheffield isolates were spread throughout the M1_UK_ phylogeny without evidence of expansion of a specific sub-clade. Two scarlet fever throat isolates did not possess *speA* consistent with lack of the *speA*-carrying Φ5005.1 phage. All other isolates had *speA*, the chromosomal *speG*, *speJ* and *smeZ*, and a combination of other prophage-associated superantigen genes. No clear correlation was seen between clinical presentation and the presence of a particular profile of superantigens and/or DNases (**Supplementary Figure 5**).

**Figure 4:**
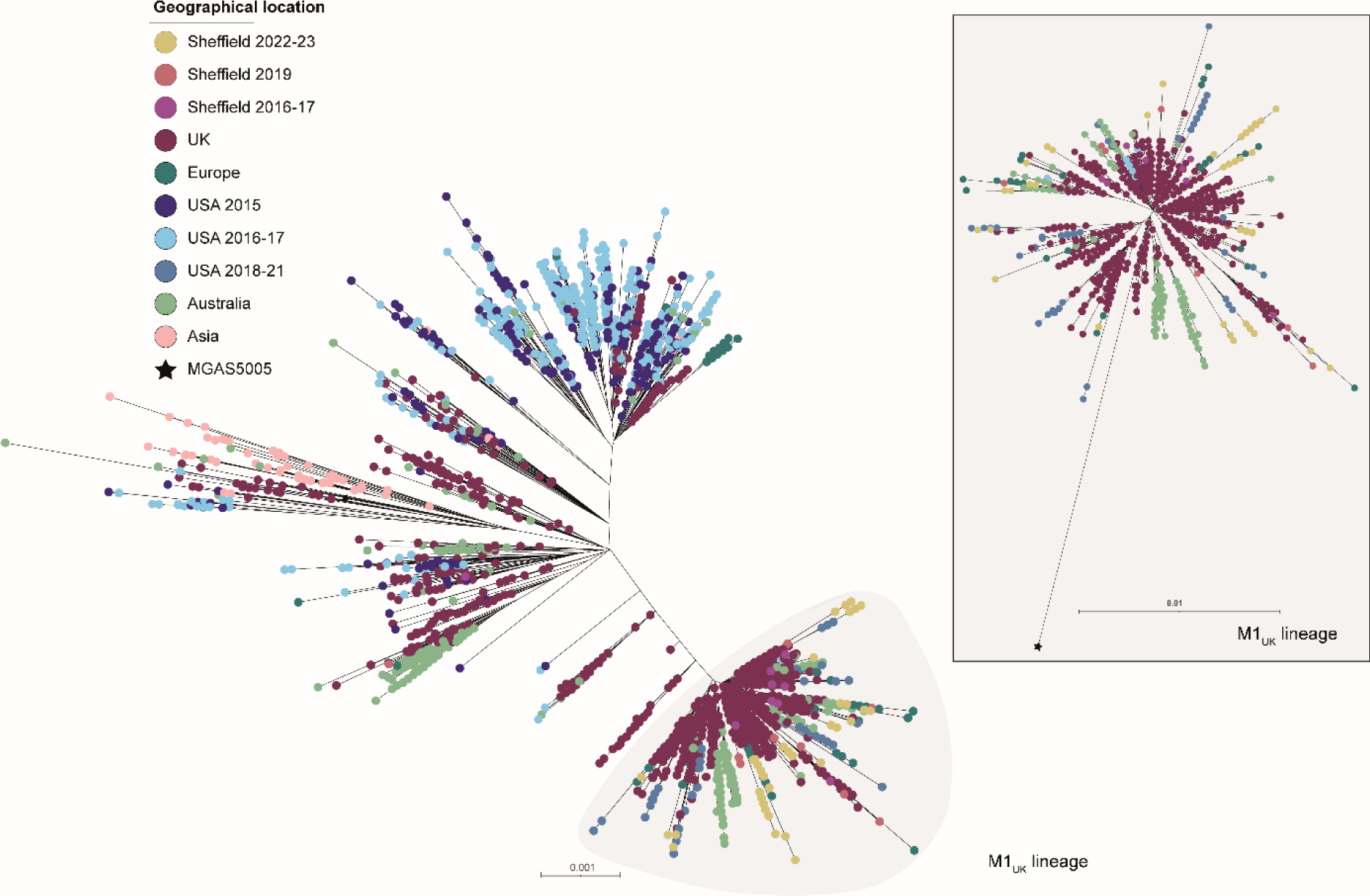
Phylogenetic analysis of Sheffield *emm*1 genomes collected in 2022-23 and 2016-17, within the context of global *emm*1 isolates. A maximum likelihood phylogenetic tree was generated with SNPs extracted from a core gene alignment (excluding prophage and SpyCI regions) to reference *emm*1 strain MGAS5005 [37]. Alongside our Sheffield *emm*1 genomes from 2022-23 and 2016-17 we included publicly available *emm*1 genome data from Sheffield, 2019 (n=14) [28]; other UK sites, 2001-2018 (n=1399) [7,31,38–42]; Denmark, 2018-2023 (n=98) [14]; Netherlands, 2019 (n=15) [13]; Portugal, 2022-23 (n=30) [16]; Australia, 2005-20 (n=318) [9]; China, 2004-16 (n=64) [43,44]; USA, 2015 (n=316) [45], 2016-17 (n=468) [46] and 2018-21 (n=85) [43]. The scale bar represents the number of nucleotide substitutions per site. The grey shaded area indicates the emergent lineage M1_UK_ defined by 27 SNPs; this lineage is shown as a separate tree within the panel.

### Emm12

A total of 64/341 (18.8%) 2022-23 isolate genomes were *emm*12, the majority of which were throat isolates (52/64, 81.3%), of which 11 were associated with scarlet fever. Ten were skin swab samples and two were ear swabs. For the 2016-17 collection, 12 out of 124 (9.7%) genomes were *emm*12 and all were from a throat source.

The core genome phylogeny of global *emm*12 isolates showed four distinct clades (**Figure 5**), as described previously [47]. The majority of all Sheffield isolates were clade I or clade IV, alongside other UK and European isolates. A single 2022-23 Sheffield isolate was clade II and no 2022-23 isolates were found within clade III. Our phylogeny also showed three sub-clades within clade IV, one of which was dominated by Sheffield, other UK and European isolates, while isolates from the USA were restricted to the other two sub-clades, one of which also included Asian strains. Other USA isolates also dominated clade II.

**Figure 5:**
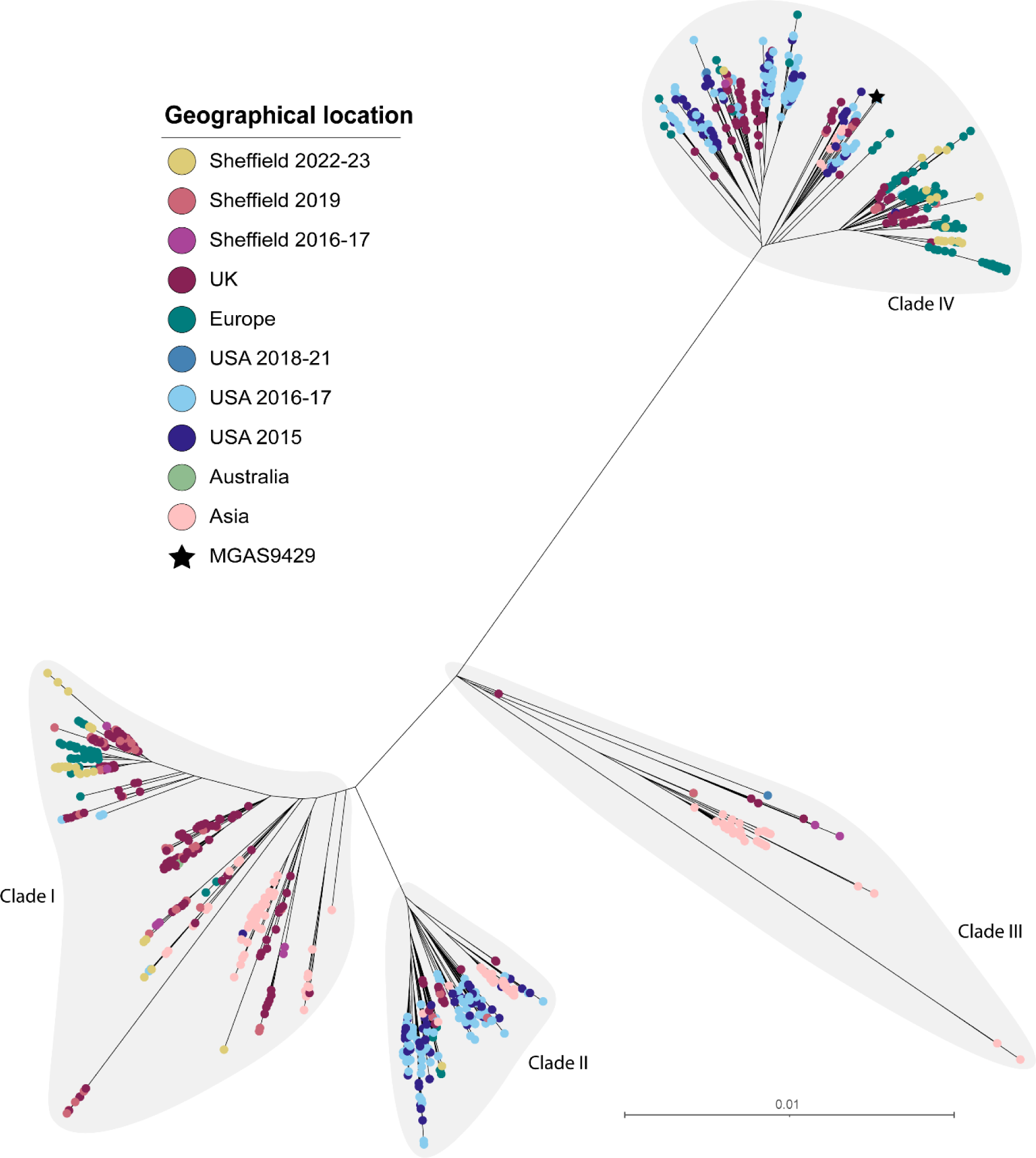
Phylogenetic analysis of Sheffield *emm*12 genomes collected in 2022-23 and 2016-17, within the context of global *emm*12 isolates. A maximum likelihood phylogenetic tree was generated with the core gene alignment (excluding prophage regions) to reference strain MGAS9429 [49]. Alongside our Sheffield *emm*12 genomes from 2022-23 and 2016-17 we included publicly available *emm*12 genome data from Sheffield, 2019 (n=15) [28]; UK, 2009-18 (n=439) [31,38,39,42]; Denmark, 2018-23 (n=239) [14]; Portugal, 2022-23 (n=12) [16]; USA, 2015 (n=134) [45], 2016-17 (n=228) [46], and 2018-21 (n=2) [11]; Australia, 1995/2017 (n=2) [47,50]; Hong Kong, 2005-11 (n=132) [47,51]; and China, 2004-18 (n= 43) [52]. Clades I-IV, previously defined by Davies et al. [47], are shaded grey. The scale bar represents the number of nucleotide substitutions per site.

Four of the 2022-23 Sheffield isolates were ST242, and clustered together within one of the sub-clades of clade IV (**Supplementary Figure 6**). All other Sheffield 2022-23 and 2016-17 isolates were ST36, except one 2022-23 isolate which was a single locus (*xpt*) variant ST1459. Seven 2022-23 throat isolates (9.1%) carried the *speA* gene and clustered together within clade I. No other 2022-23 or 2016-17 isolates carried *speA*. Again, no clear correlation was seen between clinical presentation and the presence of a particular profile of superantigens and/or DNases. Three 2022-23 throat isolates in clade I carried *mefA* and *msrD* genes, associated with acquired macrolide resistance. Interestingly, ten 2022-23 isolates with the CovR A105G variant all clustered together within clade I; a single isolate within this cluster also carried a RocA variation (G184R). A separate cluster of four isolates in clade IV carried the same mutation in RocA leading to a premature stop codon after 427 aa.

The single *emm*82 isolate from 2022-23 and all 7 *emm*82 isolates from 2016-17 had the same ST as *emm*12: ST36. These isolates were confirmed to be the lineage of *emm*82 recently identified to have arisen through recombination and *emm*-switching of *emm*12 to *emm*82 [45,48].

### Emm4

*Emm*4 was associated with invasive disease in children <15 years early in the 2022-23 upsurge [2] and was previously associated with the 2014 scarlet fever upsurge in England [4]. We found 12 *emm*4 isolates in our 2022-23 collection, of which nine were throat isolates, including three associated with scarlet fever; two were skin isolates; and one isolate was from an eye swab. One of the skin-associated isolates was ST289, highly divergent from ST39 of the other 11 strains. It was also *emm*-subtype *emm*4.2 whilst the others were *emm*4.0 or *emm*4.19, and it carried the *hasABC* locus responsible for synthesising the hyaluronic acid capsule which was characteristically absent in the other 11 *emm*4 isolates. The core-genome phylogeny also showed a highly divergent genetic background for this isolate compared to the other *emm*4 isolates.

Phylogeny of the 11 ST39 strains within a wider *emm*4 ST39 population indicated that just one strain clustered with the lineage previously described as ‘Degraded’ [53] due to substantial loss of genes within the three prophages and the integrated conjugative element (ICE) associated with *emm*4 (**Figure 6**). This lineage also has a fusion of the 5’ of *emm* gene with the 3’ of the downstream *enn* gene [54]. The five *emm*4.19 isolates clustered together within a ‘Complete’ lineage, closely related to other UK isolates. The remaining five isolates were clustered with the recently described M4_NL22_ lineage from the Netherlands, where it was associated with invasive disease [55]. This appears to be a recent emergence of this lineage in England as no *emm*4 isolates in our comparative collection from 2016-17 nor in our Sheffield 2019 isolates [28] were found clustering with M4_NL22_ isolates. Our seven Sheffield 2016-17 *emm*4 isolates included four within the ‘Degraded’ lineage whilst three were ‘Complete’. This split was similar in our 2019 Sheffield isolates with four isolates in each of these lineages. This pattern of an even divide of UK isolates between ‘Complete’ and ‘Degraded’ lineages is consistent with our phylogeny and previous findings [53], with the shift towards dominance of the ‘Complete’ lineage, at least in Sheffield, emerging during the 2022-23 upsurge.

**Figure 6:**
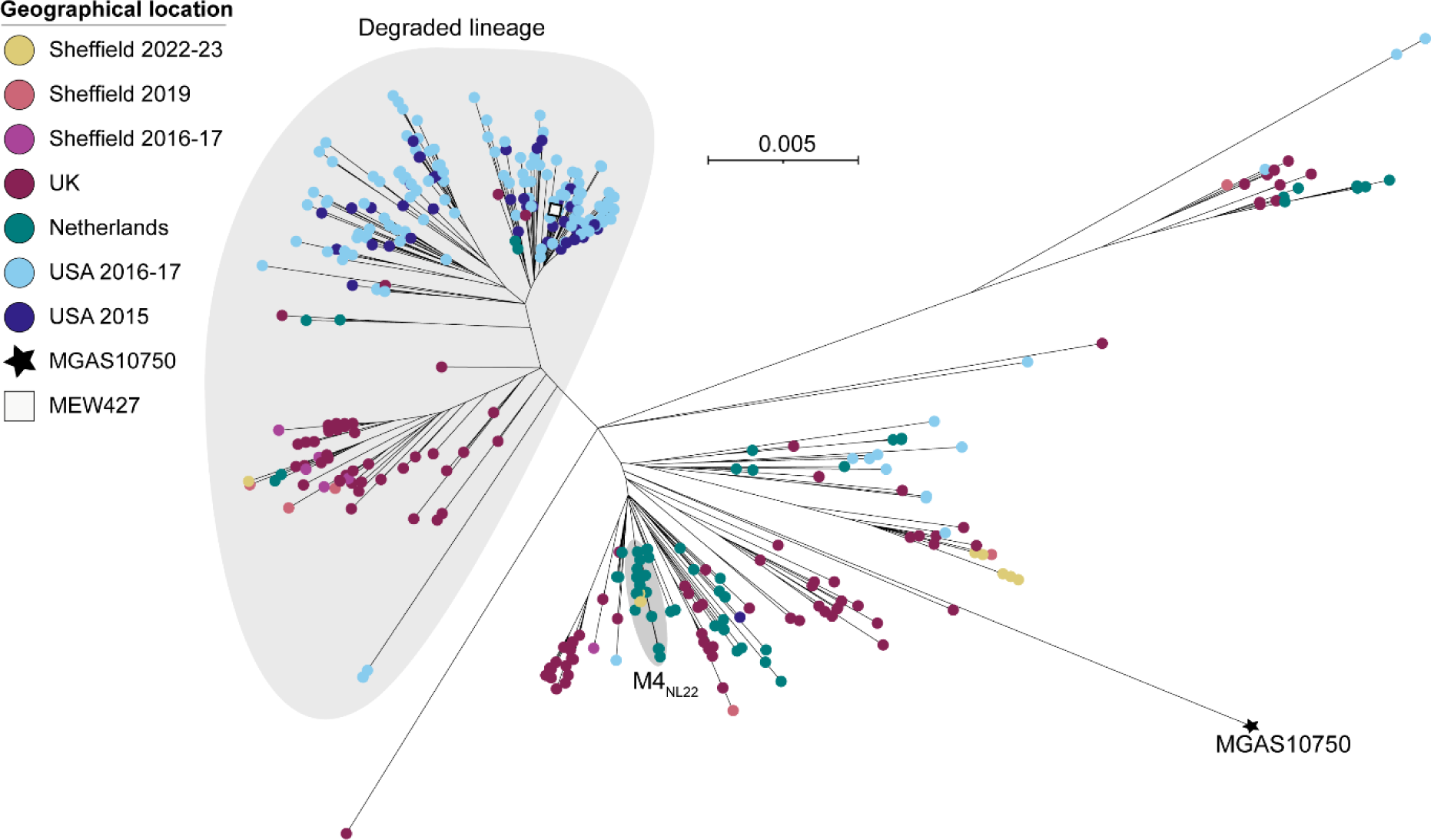
Phylogenetic analysis of Sheffield *emm*4 genomes collected in 2022-23 and 2016-17, within the context of global *emm*4 isolates. A maximum likelihood phylogenetic tree was generated with the core gene alignment (excluding prophage and SpyCI regions) to reference strain MGAS10750 (Genbank CP000262.1 [49]). The position of another reference strain MEW427 (Genbank CP014138.1 [56]) is also indicated. Alongside our Sheffield *emm*4 genomes from 2022-23 and 2016-17 we included publicly available *emm*4 genome data from Sheffield, 2019 (n=9) [28]; UK, 2011-2016 (n=176) [31, 38, 39, 42]; Netherlands, 2009-22 (n=66) [55]; and USA, 2015-17 (n=204) [45, 46]. The large grey shaded area indicates the ‘Degraded’ lineage and the small grey shaded area indicates the M4_NL22_ lineage. The scale bar represents the number of nucleotide substitutions per site.

### Emm22

A rise in *emm*22 was seen within the 2022-23 collection, overall representing 5.9% (20/341) isolates compared to none in 2016-17. Of these, four were from skin samples and the remainder from throat, with 4/16 throat samples also associated with scarlet fever. In contrast, just one isolate from our 2016-17 collection was *emm*22, from a skin source. All were ST46. Phylogenetic analysis showed the clustering of all Sheffield 2022-23 isolates, except two, together in expansion of a single lineage (**Figure 7**). All Sheffield *emm*22 isolates carried a CovR V128A variant, clustering together in a lineage with twelve other isolates from the UK, Europe and the USA all also carrying the same variant. These isolates also carried *tetM*, alongside additional strains from the same parent lineage. All *emm*22 isolates, Sheffield and globally, carried the same variation in RocA (V333A) compared to a reference RocA sequence.

**Figure 7:**
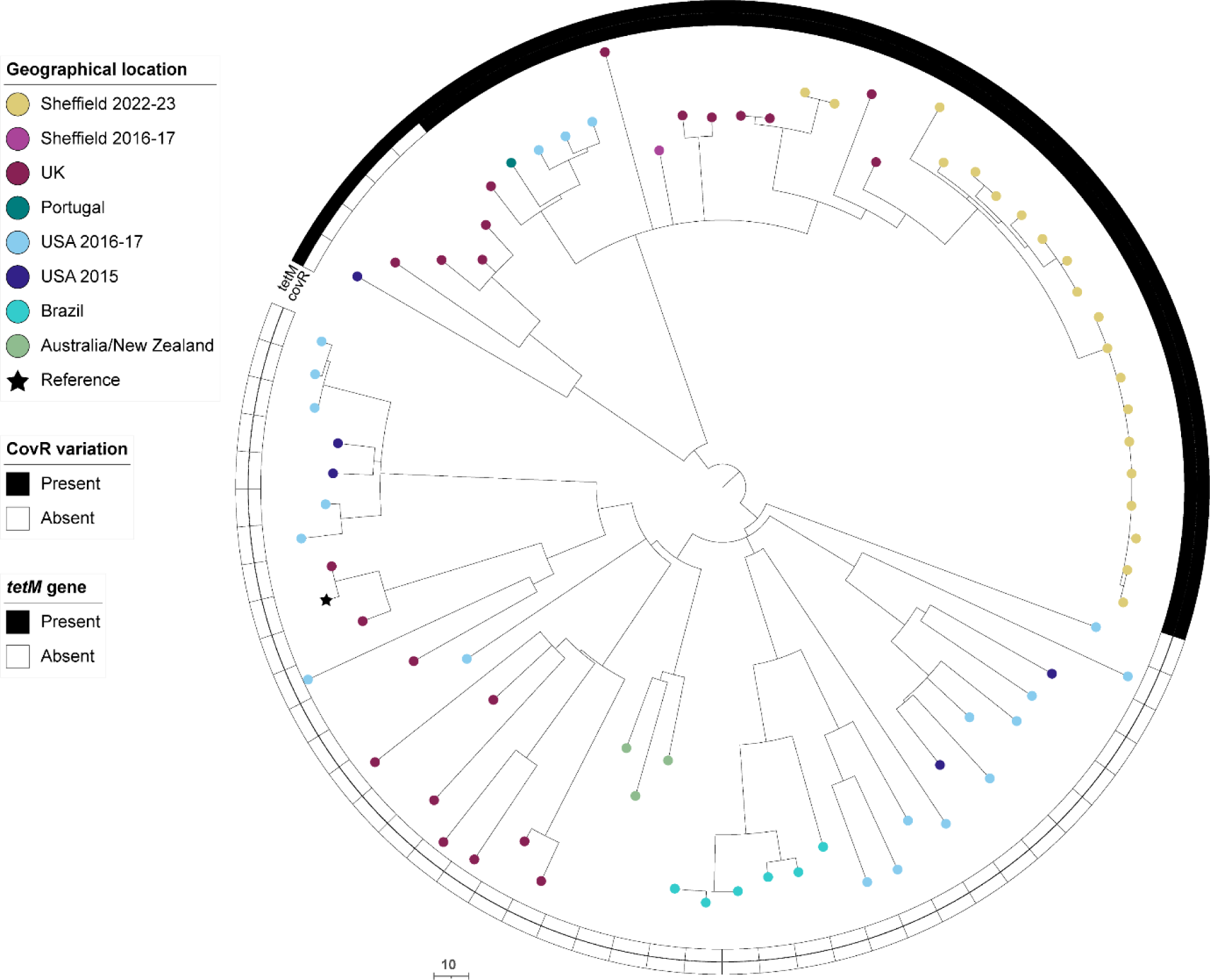
Phylogenetic analysis of Sheffield *emm*22 genomes collected in 2022-23 and 2016-17, within the context of global *emm*22 isolates. Alongside our Sheffield *emm*22 genomes from 2022-23 and 2016-17 we included publicly available *emm*22 genome data from the UK, 2001-2018 (n=22) [31, 39]; Portugal, 2022-23 (n=1) [16]; USA, 2015 (n=5) [45] and 2016-17 (n=20) [46]; Brazil, 2000-13 (n=6) [57]; Australia, 1999/2004 (n=2) [57]; and New Zealand, 2010 (n=1) [57]. A maximum likelihood phylogenetic tree was generated with the core gene alignment to the de novo assembly of the oldest UK isolate, BSAC_bs1196 [31] and removal of regions of potential recombination using Gubbins [26]. The scale bar represents the number of nucleotide substitutions per site.

### Emm75

Within all 2022-23 Sheffield isolates, 4.4% were *emm*75 (15/341), of which ten were from a throat source, four from skin and one from an ear swab. Of the ten throat isolates, four were associated with scarlet fever. By comparison, in 2016-17, 9.1% of isolates were *emm*75 (15/165), all of which were throat isolates not associated with scarlet fever. All Sheffield *emm*75 isolates from both 2022-23 and 2016-17 were ST150, distinct from the other dominant lineage of *emm*75, ST49, and all carried the same variation in *covS* (S337L). Of the 2022-23 isolates, 13/15 clustered together in a new sublineage and had identical superantigen profiles (**Figure 8**). Within this cluster, 12/13 were found to have the same additional T in a seven residue homopolymeric tract, leading to a premature stop codon after 46 amino acids, and therefore are likely to be acapsular. This is new compared to 2016-17 Sheffield isolates, none of which carried a *hasA* mutation and were scattered throughout the phylogeny. Other acapsular *emm*75 strains with the same premature stop codon in HasA were identified in a small cluster within USA isolates, and a single sporadic UK strain.

**Figure 8:**
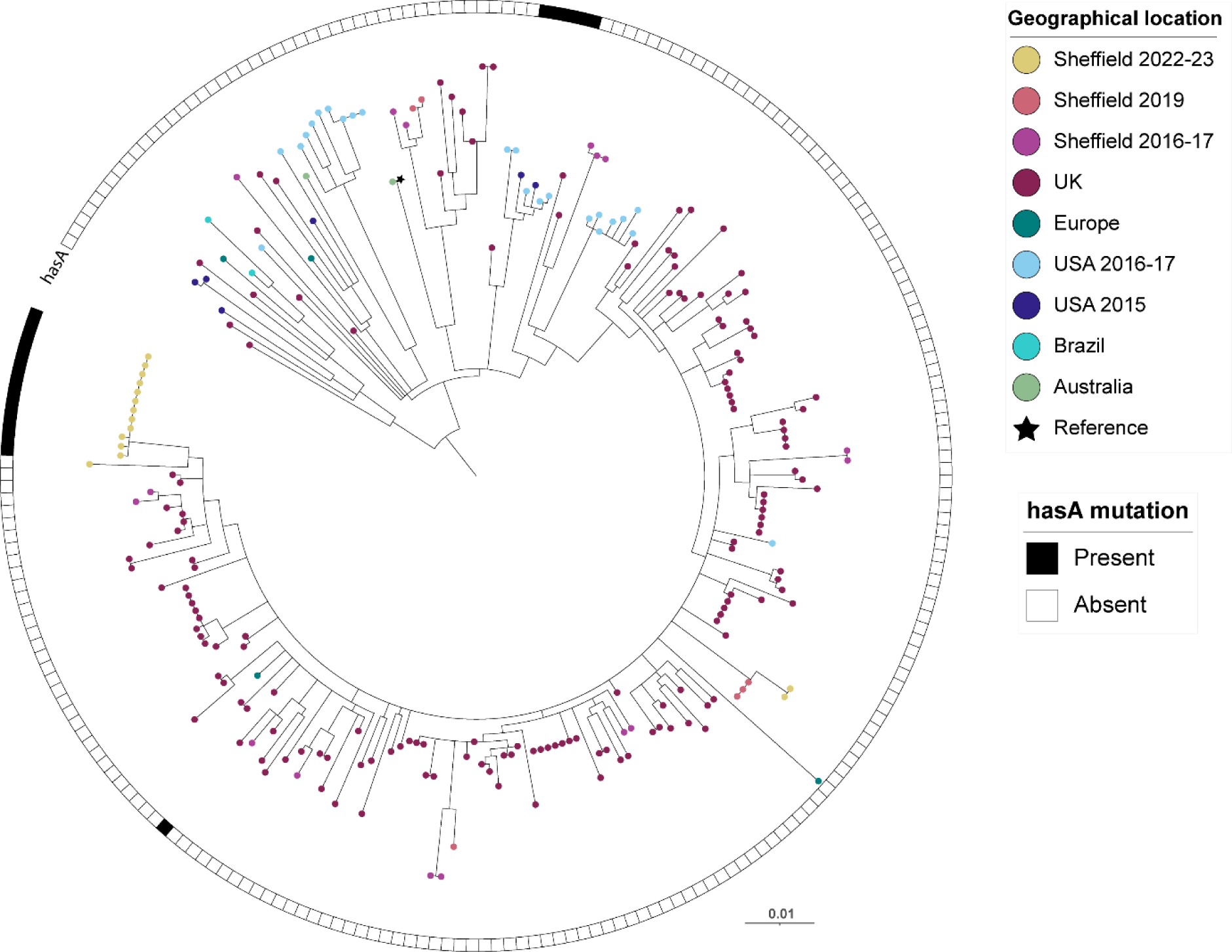
Phylogenetic analysis of Sheffield *emm*75 genomes collected in 2022-23 and 2016-17, within the context of global *emm*75 isolates. A maximum likelihood phylogenetic tree was generated with the core gene alignment to reference strain M75 (star) (Genbank CP033621.1 [58]). Alongside our Sheffield *emm*75 genomes from 2022-23 and 2016-17, we included publicly available *emm*75 genome data from Sheffield, 2019 (n=6) [28]; UK, 2011-16 (n=158) [31, 38, 39, 42]; Belgium, 2004 (n=1) [57]; USA, 2015-17 (n=81) [45, 46]; Australia, 1991-2010 (n=7) [57]; and Brazil, 2007-12 (n=2) [57]. The scale bar represents the number of nucleotide substitutions per site. The presence or absence of a *hasA* nonsense mutation is indicated for each strain. Only isolates of ST150 (and closely related STs) are shown, excluding the other major lineage of *emm*75 (ST49 and closely related) as these are genetically quite distant [31]. Those that were ST49, and closely related STs comprised genomes from UK, 2014-18 (n=3) [39]; USA, 2015 (n=14) and 2016-17 (n=38) [45, 46]; Australia, 1991-04 (n=6) [57]; Fiji, 2006 (n=3) [57]; India, 2007-2010 (n=2) [57]; and Kenya, 2000-11 (n=5) [57]. No Sheffield isolates were ST49.

### *Emm*76, 77 and 87

Previously we found recent expanding lineages within *emm*76, 77 and 87 that were acapsular due to mutations in *hasA,* and had undergone recombination around the *nga-ifs-slo* promoter region to become high-toxin expressing variants [31]. In our 2022-23 isolates this was still the case for *emm*77 and *emm*87 but not for *emm*76.

In 2022-23, 5.6% (19/341) of isolates were *emm*76, compared to none in 2016-17. Of these, 18 were from a skin source and one from an ear swab and all were ST378, predicted to be encapsulated as they had no mutations in *hasA* or *hasB*, with low-toxin expression (**Supplementary Figure 7**). All carried *tetM*, as described previously within this lineage [31]. This reflects a recent expansion of the encapsulated, low-toxin phenotype ST378 lineage, in contrast to the previously-described ST50 lineage which was acapsular with high-toxin expression [31].

In 2022-23, 4.1% (14/341) of isolates were *emm*77, an increase from 2.4% (4/165) in 2016-17. Of the 2022-23 isolates, 8 were from throat swabs and not associated with scarlet fever, 5 were from a skin source and one from an ear swab. The majority (11/14) were ST63 and part of the previously described acapsular (truncated HasA after 154 aa) with high-toxin expression lineage [31]. Isolates in this lineage also carry *tetO* and *ermA* as well as the CovR M170I variant. The two other *emm*77 isolates were ST399, quite distinct from ST63 although also predicted to be acapsular (truncated HasA after 46 aa) but with low-toxin expression. These ST399 isolates carried *tetM* and had a T266I variation in CovS. An increase in *emm*87 was also observed in 2022-23 to 6.7% (23/341) from 4.8% in 2016-17. Of the 2022-23 isolates, 16 were from a throat source and two from ear swabs, two from eye swabs, and three from skin sources. All isolates were part of the previously described lineage characterised as being acapsular, through a mutation in *hasA* (truncating HasA after 46 aa), and high-toxin expressing through recombination around the *nga-ifs-slo* locus and promoter [31] (**Supplementary Figure 8**). Interestingly, international phylogeny of *emm*87 revealed four broad clades, one of which contains a sub-lineage dominated by Sheffield and other UK strains (**Supplementary Figure 8**). This sub-lineage is characterised by the same mutation in *hasB* resulting in a premature stop codon after 188 aa, in addition to the mutation in *hasA*. Other Sheffield isolates found elsewhere within the phylogeny did not carry any mutations in *hasB*, though small clusters of US *emm*87 isolates carried other truncating mutations in *hasB*. There was also some evidence of geographical variation across the lineages, with USA isolates dominating one lineage and UK isolates another.

## Discussion

In recent years the UK has experienced significant rises in morbidity and mortality in association with substantial upsurges in *S. pyogenes* infections, including in 2017-18 and in 2022-23 [3]. Despite this, study of non-invasive isolates has been limited and the capacity for dynamic changes in non-invasive disease to drive upsurges and invasive disease is poorly understood. We sequenced 341 non-invasive isolates from Sheffield during the major 2022-23 UK upsurge and although we found *emm*1 and *emm*12 to be the leading causes of both throat and skin infections, they were differentially followed by *emm*22, *emm*87, and *emm*89 in throat infections but *emm*76 and *emm*49 in skin. A comparison to non-upsurge isolates from 2016-17, indicated *emm*1 and *emm*12 contributed significantly to the 2022-23 upsurge but other *emm*-types had also changed over time with more *emm*22 in throat infections and *emm*76 in skin infections. All 2022-23 *emm*1 isolates were the prevalent M1_UK_ lineage but more diverse lineages were identified in other *emm*-types, including *emm*12, and emergent lineages in others, including *emm*75, demonstrating that, at least local to Sheffield, the upsurge was not primarily caused by a single genotype.

For both time periods studied, non-invasive *S. pyogenes* infections were most common in those aged 0-9 years, but more infections were seen in 5-9 year old children in 2022-23 than 2016-17 (33.3% vs 15.7%). Overall, those aged 4, 5, 6 or 7 years of age had the highest rates of *S. pyogenes* infection of any age in 2022-23. This may reflect accelerated exposure to infection associated with school attendance, in keeping with reduced immunity to *S. pyogenes* and common respiratory viruses in this age group following reduced exposure during the COVID-19 pandemic [2, 59].

The dominant peak in throat isolates at week 49 of 2022 within our collection, followed by a second smaller peak in week 3 of 2023, coincides with the peaks in scarlet fever notifications reported by UKHSA during this time period [3]. During week 49, NHS England group A *Streptococcus* interim clinical guidance summary for case management was issued with guidance temporarily altering the clinical scoring criteria threshold for immediate antibiotic treatment in children with a sore throat. The issue of this guidance on 9th December 2022 likely enhanced clinician confidence in the empirical diagnosis of *S. pyogenes* throat infection and therefore resulted in a fall in throat swabs being received by the diagnostic laboratory. In contrast, the prominent peak in skin isolates occurred in week 52, between the peaks in throat isolates. This may be due to behavioural or environmental factors, such as increased skin-to-skin transmission of *S. pyogenes* associated with increased social mixing or altered chronic wound care at this time of year.

During the time of our 2022-23 collection, the Department of Laboratory Medicine, Sheffield, identified 19 iGAS samples (*S. pyogenes* isolated from a sterile site) and these had undergone routine *emm*-typing by UKHSA, but genomic data was not available. Patients ranged in age from 1 to 94 years, but were most commonly 0-9 years (36.8%, 7/19) or over 75 years (31.6%, 6/19), in keeping with national data from the same time period. By far the majority were *emm*1 (57.9%, 11/19), followed by *emm*76 (15.8%, 3/19), *emm*89 (10.5%, 2/19), *emm*92 (10.5%, 2/19), and *emm*87 (5.3%, 1/19). The most common sample type was blood culture, at 13/19 samples (68.4%, 3/19), followed by empyema samples (15.8%, 3/19), and soft tissue or joint fluid (15.8%, 3/19). All empyema samples were *emm*1 and all in children aged 0-9 years. Analysis of national data from this time period has identified a significant association between M1_UK_ and pleural isolates, likely the result of an early *S. pyogenes* upsurge coinciding with the respiratory virus season, facilitating disease progression [60]. Nationally, *emm*1 and *emm*12 were the most common *emm*-types amongst invasive isolates in all age groups during this upsurge, notably with *emm*4 as the next most common *emm*-type in children [2]. Whilst we identified that 57.9% of our invasive isolates were *emm*1, our dataset did not identify any invasive *emm*12 nor *emm*4 isolates and the remaining 47.4% (8/19) of invasive isolates during the 2022-23 collection were of four other *emm*-types, potentially a reflection of regional variation in circulating strains within England. The skin prevalent *emm*76 did cause a substantial proportion of skin and iGAS infections suggesting we should not overlook this infection site. It is not known if the pronounced association of *emm*76 with skin infections was a local phenomenon as national data on skin infection *emm*-types is not collected.

Tissue tropism within *S. pyogenes* infections is well-recognised but bacterial molecular adaptations associated with tissue specialisation remain incompletely understood [61]. An acapsular genotype was frequently seen in throat isolates in this study, with just 51% predicted to be able to produce capsule compared to 78% of skin isolates. In throat isolates, these acapsular strains were predominantly ‘generalist’ pattern E. Just 27% of pattern E isolates overall were predicted to be encapsulated and made up 44.5% of throat infections. Interestingly, although more skin than throat infections were pattern E, at 56%, a higher proportion (62.5%) of these were encapsulated. We also identified evidence of evolving capsule loss with the emergence of a recent acapsular sublineage of *emm*75, a pattern E type, and the increase in prevalence of acapsular *emm*22.

We further observed tissue-specific differences in predicted expression of the toxins NADase and SLO based on the promoter sequence, with 90% of throat isolates predicted to express high levels of toxin compared to 56% of skin isolates, highlighting a key role for toxin expression in the pathogenesis of *S. pyogenes* throat infections. We previously provided evidence of convergent evolution with acapsular strains gaining increased toxin expression through homologous recombination [31]. Successful emergent lineages characterised by this recombination event have been identified previously in pattern E *emm*76, *emm*77, and *emm*87. Interestingly, whilst we continued to see expansion of these acapsular/high-toxin lineages in *emm*77 and *emm*87, in *emm*76 we observed instead an expansion of an ST378 encapsulated/low-toxin lineage to become a leading cause of skin infections. Indeed, overall we observed significant differences between toxin/capsule genotypes by infection site, with more throat isolates possessing a high-toxin/acapsular genotype and more skin isolates possessing a low-toxin/encapsulated genotype. An association between NADase activity and tissue tropism has also been identified previously, with NADase-inactive strains being primarily ‘skin-associated’ *emm*-pattern D, suggesting toxin may be less essential for *S. pyogenes* infection in the skin ecological niche compared to in the throat [62].

The most frequently identified *emm*-type across both throat and skin infections in 2022-23 was *emm*1; all were M1_UK_ lineage and distributed throughout the M1_UK_ phylogeny without evidence of local expansion of a specific sub-clade. In contrast, *emm*12 isolates from our collections in both 2022-23 and 2016-17 were distributed predominantly across two of the four clades: I and IV. Previous work has suggested an association in clades I-III between scarlet fever and *ssa* (encoding streptococcal superantigen A), and an absence of scarlet fever in clade IV, however no such clear associations were seen in our collection [47]. Geographical variation was apparent across our *emm*12 phylogeny, with the majority of Sheffield isolates found within a single subclade of clade IV and clade I, in a distribution distinct from that of US isolates and again from Asian isolates. We observed similar geographical variation in *emm*87, with two of the four main clades dominated by USA strains and one clade containing the majority of Sheffield and other UK strains. These variations are in keeping with regional divergence of circulating lineages and can give rise to the emergence of geographically-restricted sublineages, some of which have been seen to expand more widely if carrying a fitness advantage [7].

Overall, we found that an increase in prevalence of *emm*1 and *emm*12 in non-invasive disease in 2022-23 locally reflected the national increase of these *emm*-types in iGAS and scarlet fever cases. We have highlighted the need to study both throat and skin infection isolates as they can differ, and the potential for differential capsule expression and/or NADase and SLO toxin expression to drive tissue tropism. We also demonstrated that *emm*-types may not be represented by single lineages and these can change over time and by geographical location. Frequent monitoring with WGS is needed to determine how rapidly these changes occur, what factors influence these changes, and how they might drive infection rates, including overspill from non-invasive disease into invasive and upsurges.

## Acknowledgements

The authors are grateful to laboratory colleagues at Sheffield Teaching Hospitals NHS Foundation Trust for isolate collection. We also acknowledge the input of the genomics pipeline group at the Earlham Institute. CET was a Royal Society & Wellcome Trust Sir Henry Dale Research Fellow (208765/Z/17/Z) and then a Wellcome Trust Career Development award holder (227240/Z/23/Z).

## Supplementary

**Supplementary Figure 1:**
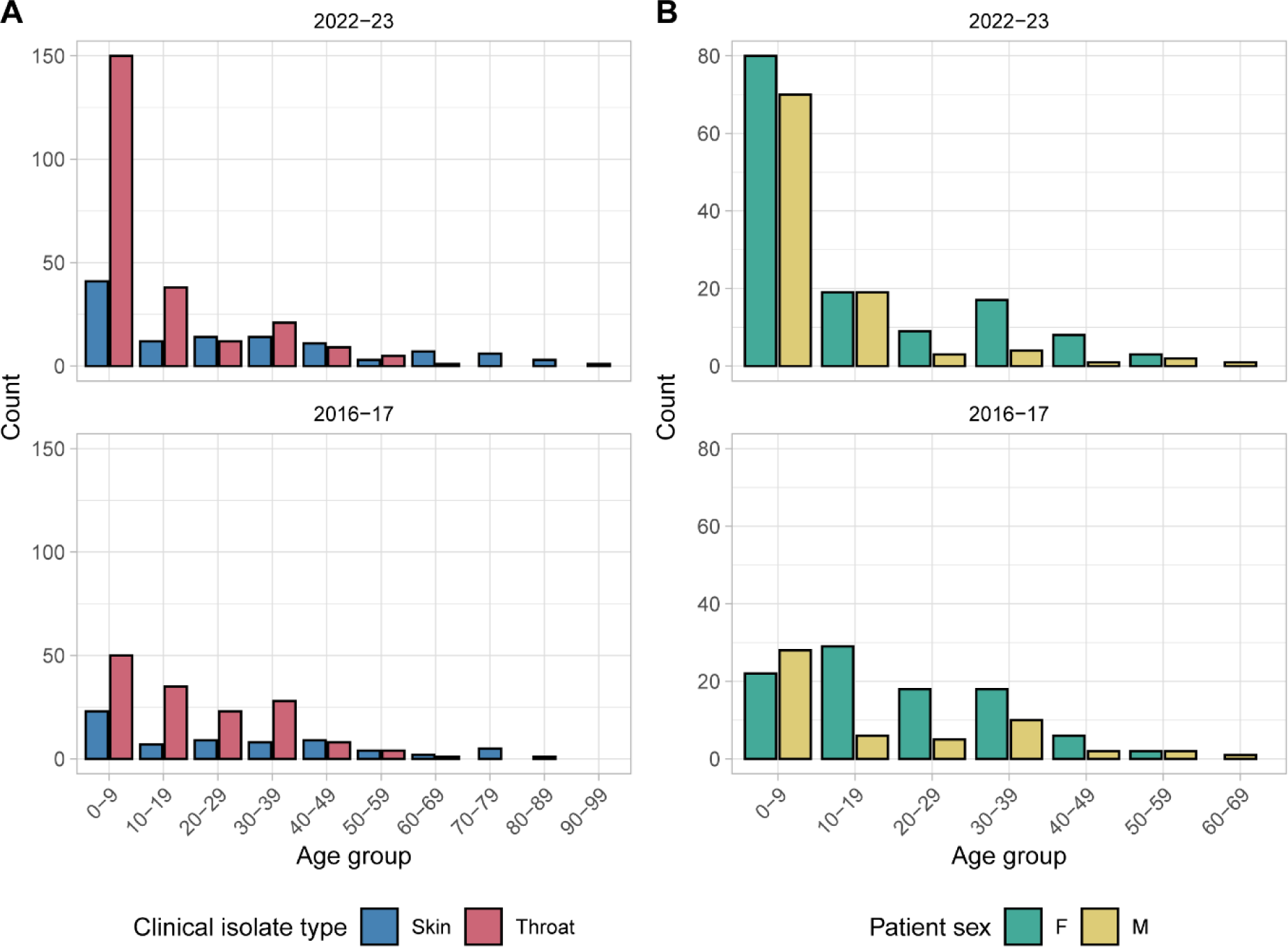
(**A**) Age distribution of patients from whom isolates were collected in 2022-23 and 2016-17, grouped by clinical isolate type. Other clinical isolate types (eye, ear and nose) are excluded. (**B**) Age distribution of patients across throat isolates in 2022-23 and 2016-17, grouped by patient sex.

**Supplementary Figure 2:**
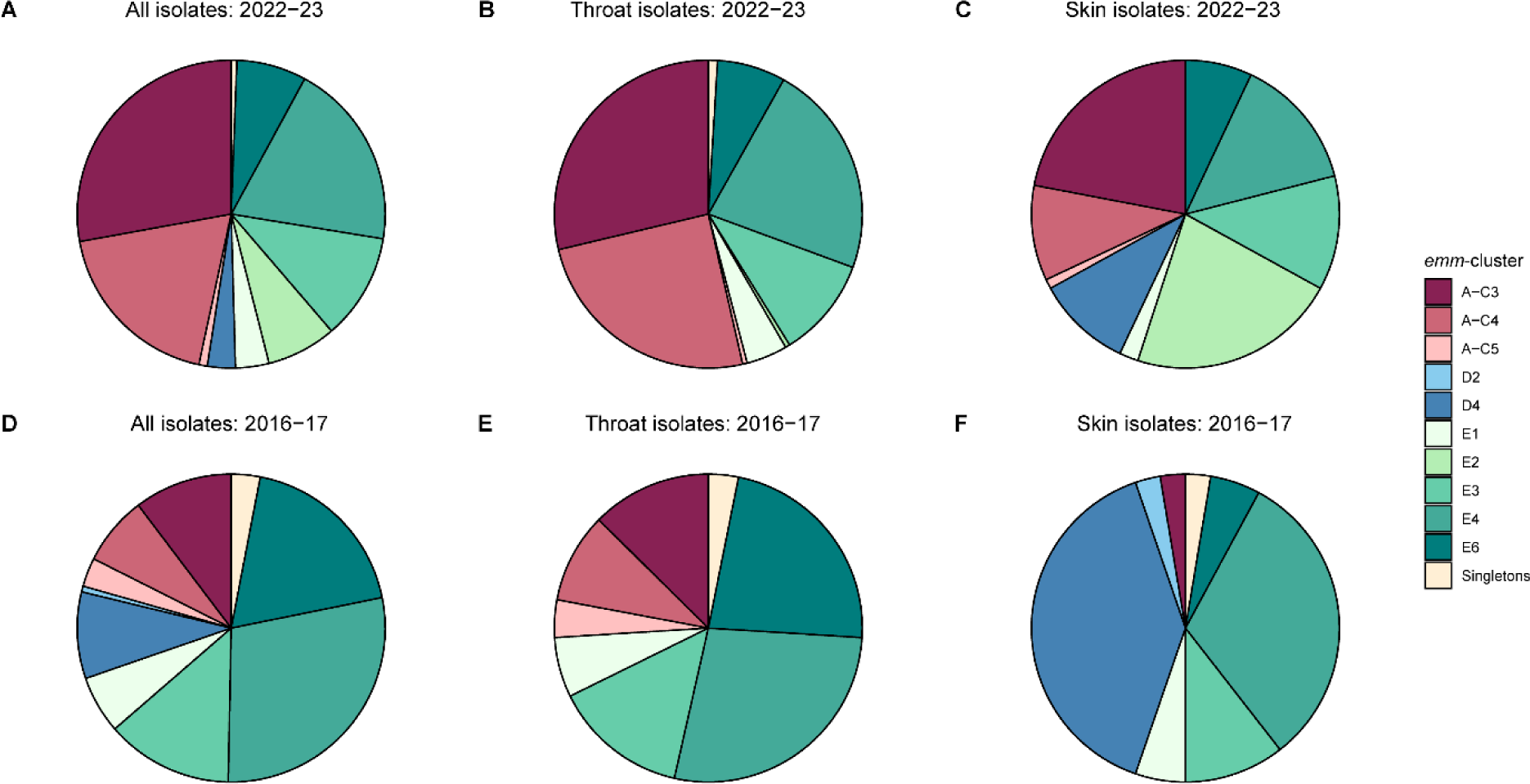
Distribution of *emm*-clusters across 2022-23 and 2016-17 collections. (**A**) All 2022-23 isolates; (**B**) 2022-23 throat isolates; (**C**) 2022-23 skin isolates; (**D**) all 2016-17 isolates; (**E**) 2016-17 throat isolates; (**F**) 2016-17 skin isolates. Pie charts represent the percentage of isolates associated with each cluster.

**Supplementary Figure 3:**
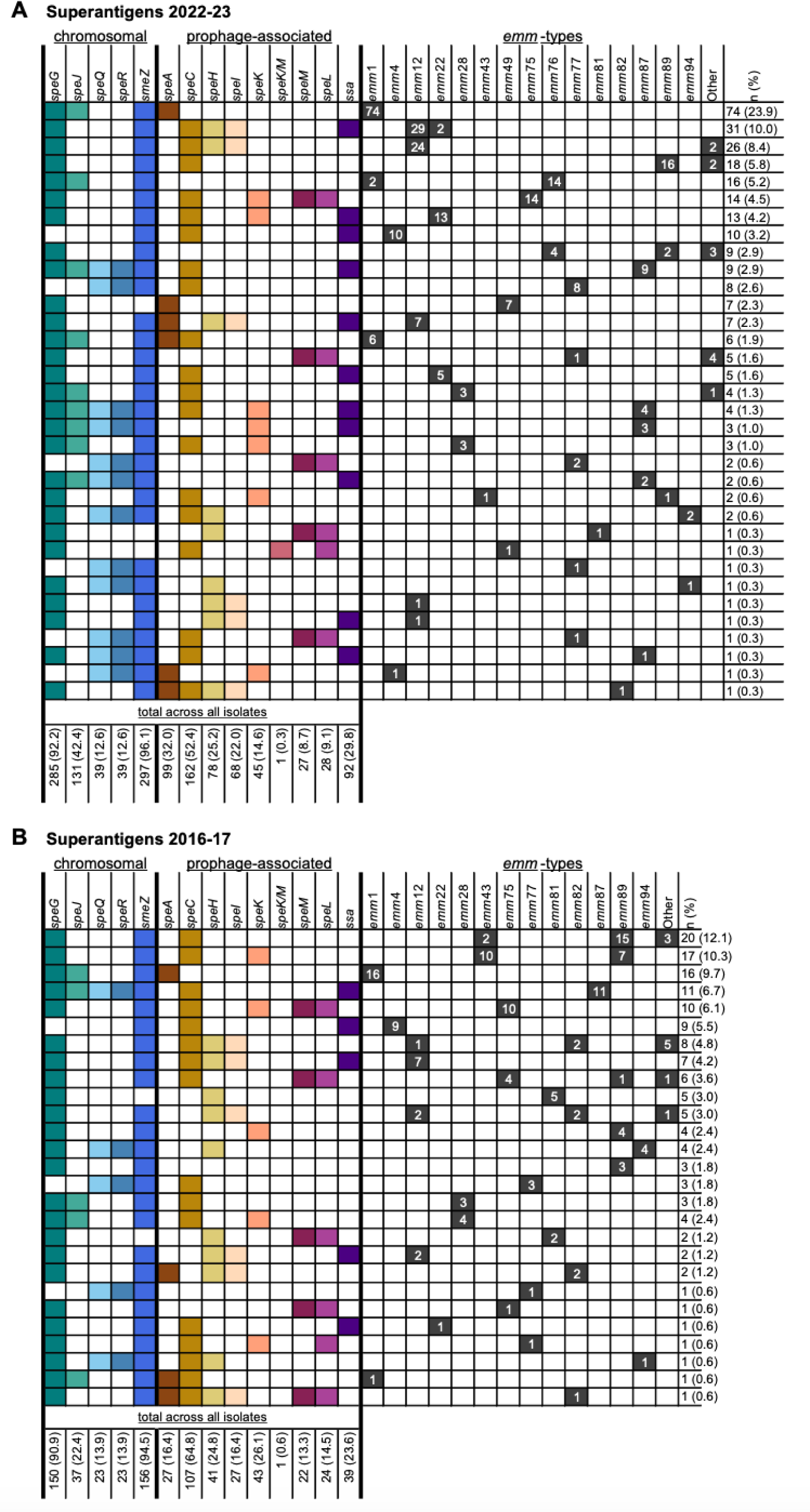
Summary of superantigen profiles by *emm*-type in throat and skin isolates collected in (A) 2022-23 and (B) 2016-17. Data shown for the most common *emm*-types ordered from most common profile to least common; other *emm*-types are categorised here as ‘Other’. Superantigen combinations found only in ‘Other’ *emm*-types have been excluded. Individual superantigen totals at the bottom of the figure are displayed as n (%) and represent all skin and throat isolates of all *emm*-types, including those only found in ‘other’ *emm*-types.

**Supplementary Figure 4:**
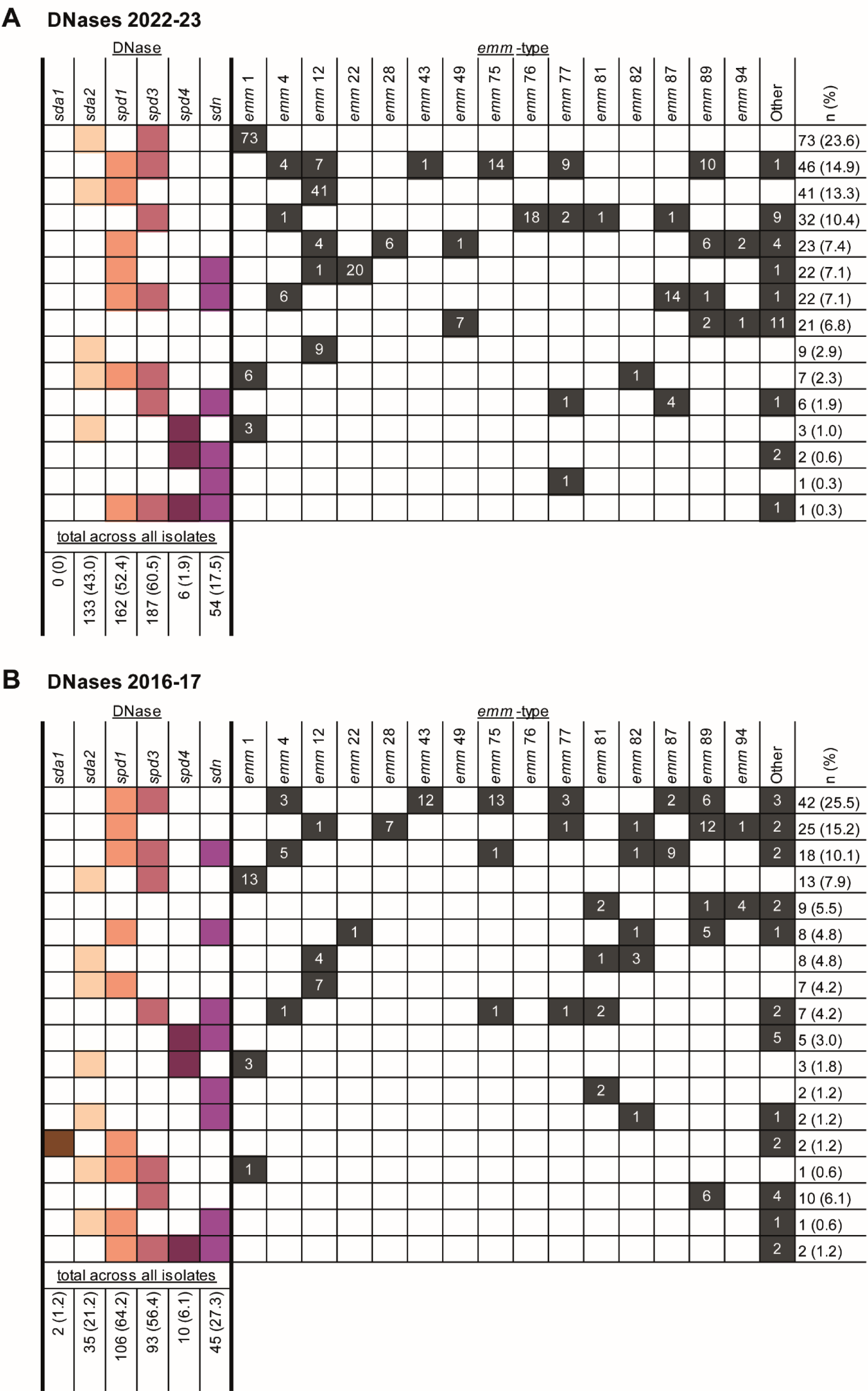
Summary of DNase profiles by *emm*-type in throat and skin isolates collected in (A) 2022-23 and (B) 2016-17. Data shown for the most common *emm*-types ordered from most common profile to least common; other *emm*-types are categorised here as ‘Other’. Individual DNase totals at the bottom of the figure are displayed as n (%) and represent all skin and throat isolates of all *emm*-types.

**Supplementary Figure 5:**
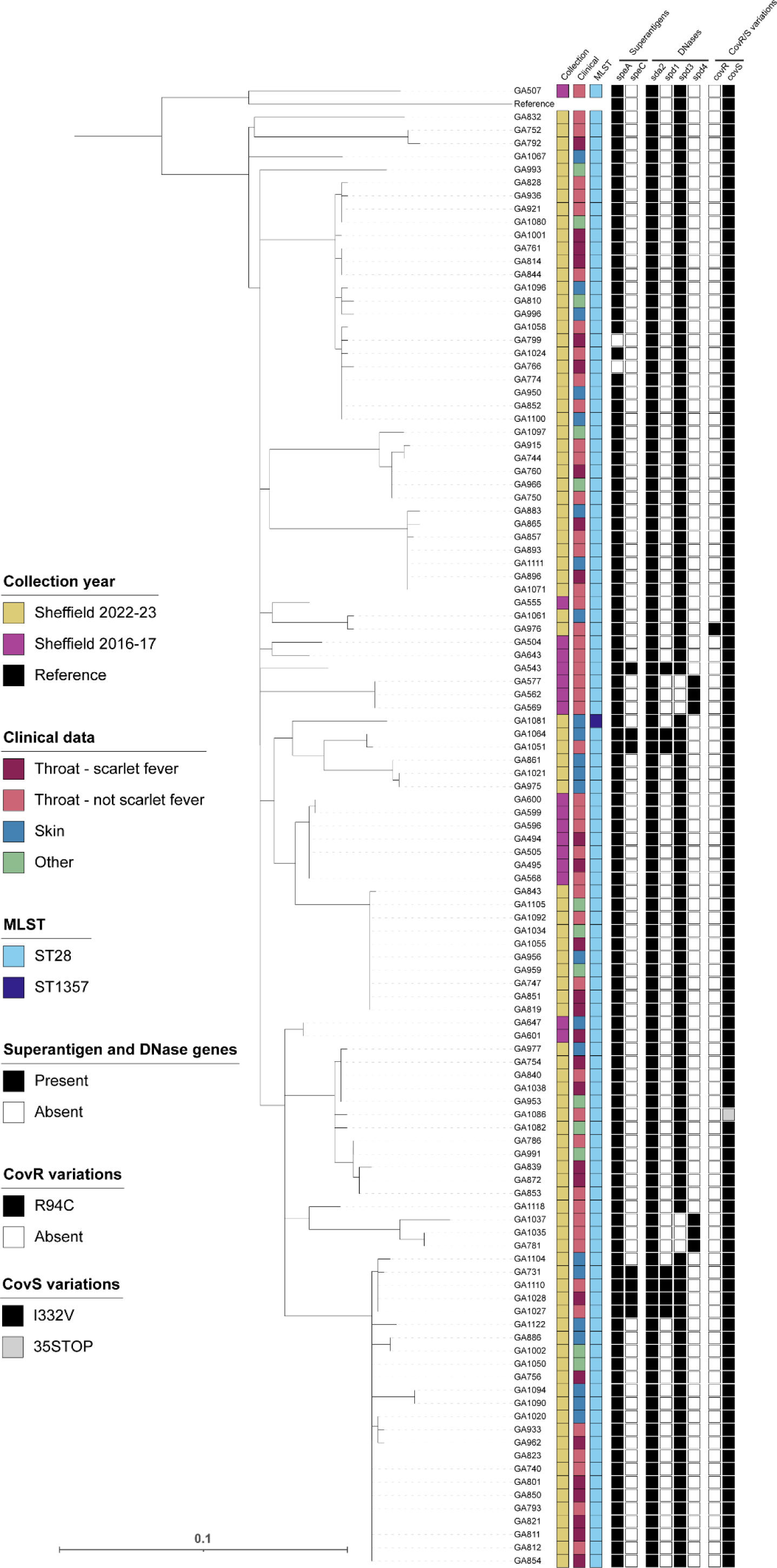
Phylogenetic analysis of the 112 Sheffield *emm*1 genomes collected in 2022-23 and 2016-17. A maximum likelihood phylogenetic tree was generated with the core gene alignment to reference strain MGAS5005. The scale bar represents the number of nucleotide substitutions per site. Colour strips indicate the year of isolate collection, clinical presentation and the multi-locus sequence type (MLST). The presence (black) and absence (white) of superantigen genes *speA* and *speC*; prophage-associated DNase genes *sda2*, *spd1*, *spd3*, and *spd4* are indicated. Variations in CovR and CovS are indicated. All isolates possessed chromosomal superantigens *speG*, *speJ* and *smeZ*. No isolates had superantigen genes *speH*, *speI*, *speK*, *speL*, *speK/M*, *speM*, *speQ*, *speR* or *ssa*, nor DNase genes *sda1* or *sdn*. No antimicrobial resistance (AMR) genes were identified within the Sheffield *emm*1 isolates. All Sheffield *emm*1 isolates also had a Q259R variation in RocA.

**Supplementary Figure 6:**
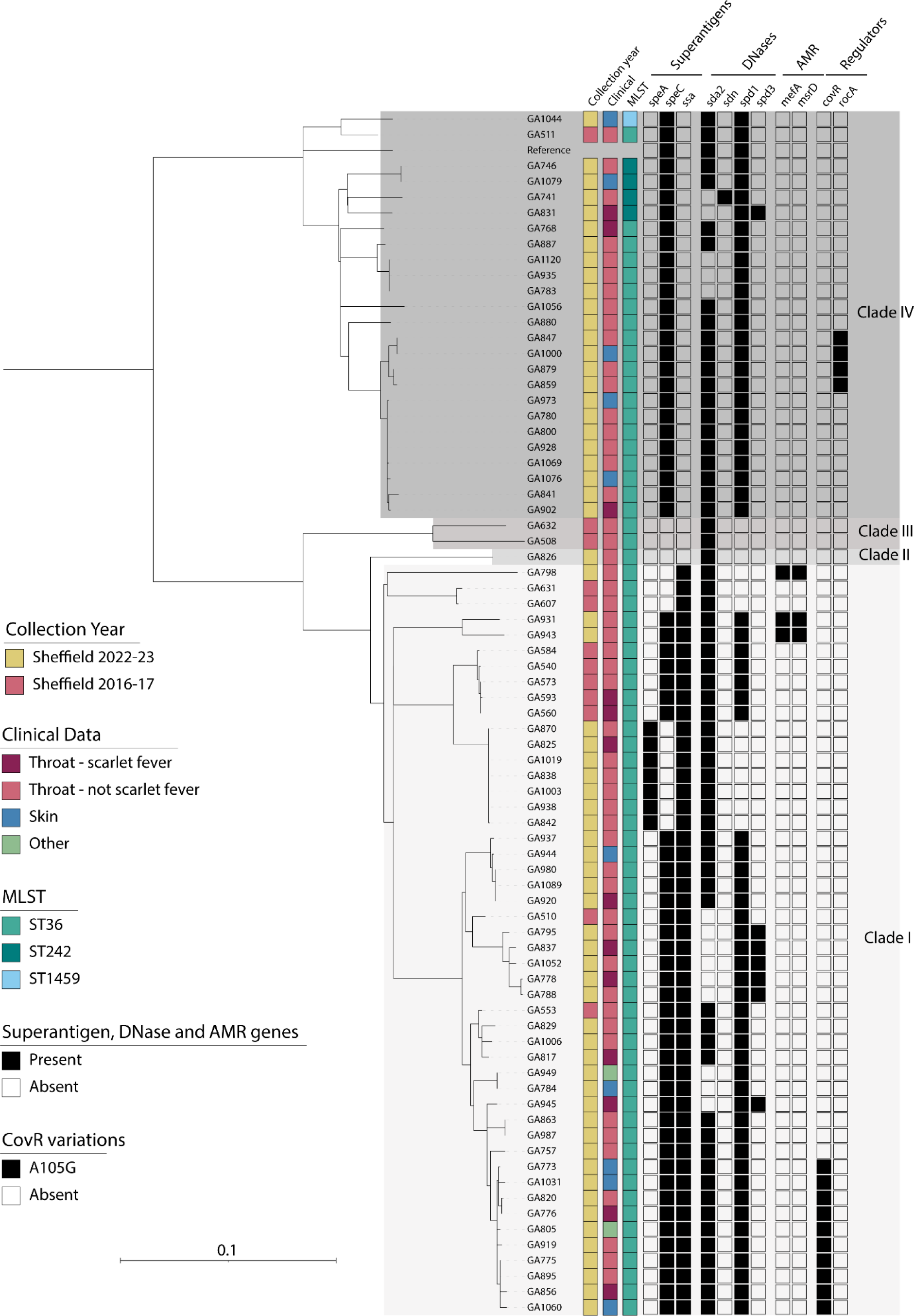
Phylogenetic analysis of the 77 Sheffield *emm*12 genomes collected in 2022-23 and 2016-17. A maximum likelihood phylogenetic tree was generated with the core gene alignment to reference strain MGAS9429. The scale bar represents the number of nucleotide substitutions per site. Colour strips indicate the year of isolate collection and the multi-locus sequence type (MLST). The presence (grey) and absence (white) of superantigen genes *speA*, *speC* and *ssa*; DNase genes *sda2*, *sdn, spd1* and *spd3*; and variations in CovR and RocA are indicated. No variations in CovS were identified within Sheffield *emm*12 genomes. All isolates possessed DNase genes *speG, speH* and *speI*. No isolates had superantigens *speJ, speK, speK/M, speM, speL, speQ, speR* or *smeZ*, nor DNase genes *sda1* and *spd4*. The presence (grey) and absence (white) of antimicrobial resistance (AMR) genes *mefA* and *msrD* is indicated; no other AMR genes were identified within Sheffield *emm*12 isolates.

**Supplementary Figure 7:**
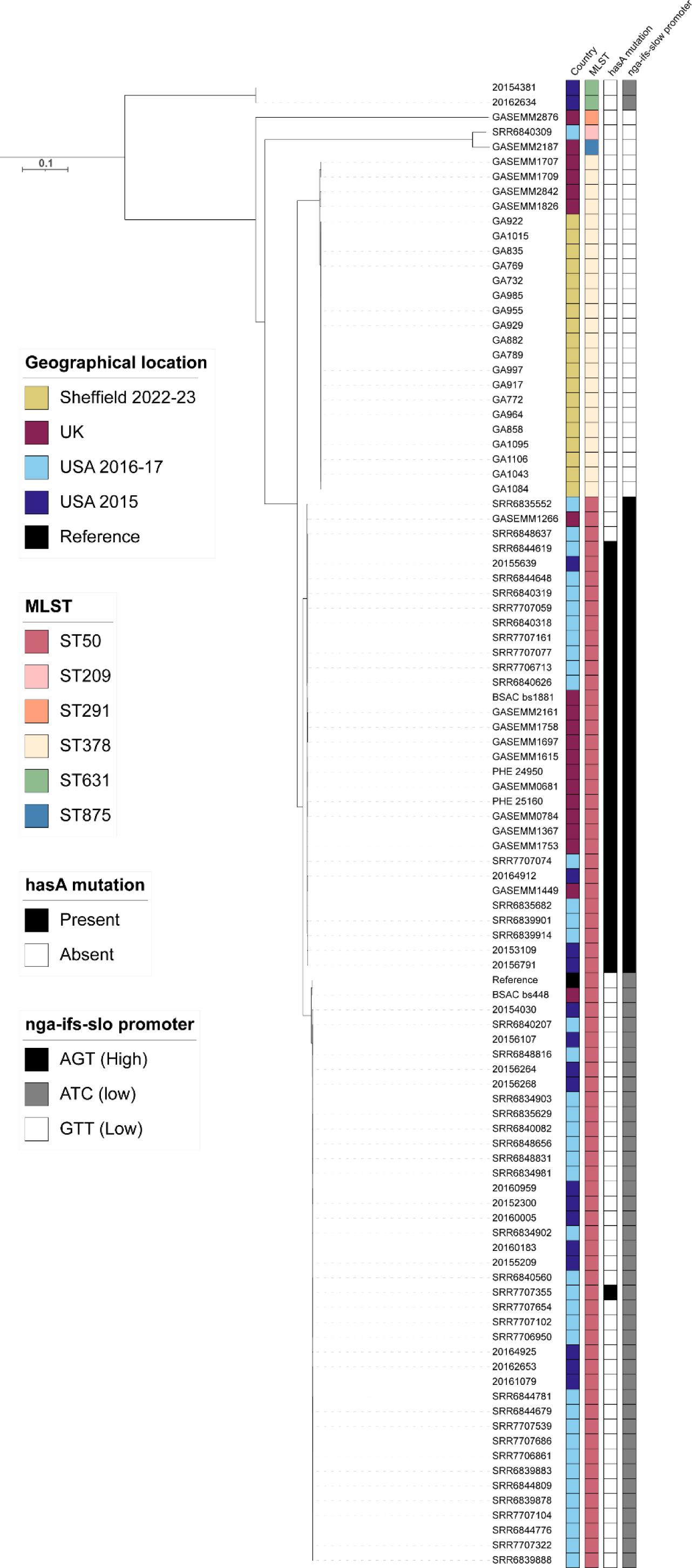
Phylogenetic analysis of Sheffield *emm*76 genomes collected in 2022-23, within the context of other UK and USA *emm*76 isolates. Alongside our Sheffield *emm*76 genomes from 2022-23 and 2016-17 we included publicly available *emm*76 genome data from the UK, 2002-2018 (n=20) [31,38,39]; USA, 2015 (n=18) [45] and 2016-17 (n=42) [46]. A maximum likelihood phylogenetic tree was generated with the core gene alignment to reference strain BSAC_bs448. The scale bar represents the number of nucleotide substitutions per site. Colour strips indicate the country of isolate collection and the multi-locus sequence type (MLST). The presence (black) and absence (white) of *hasA* mutations is indicated, as are variants of the *nga-ifs-slo* promoter.

**Supplementary Figure 8:**
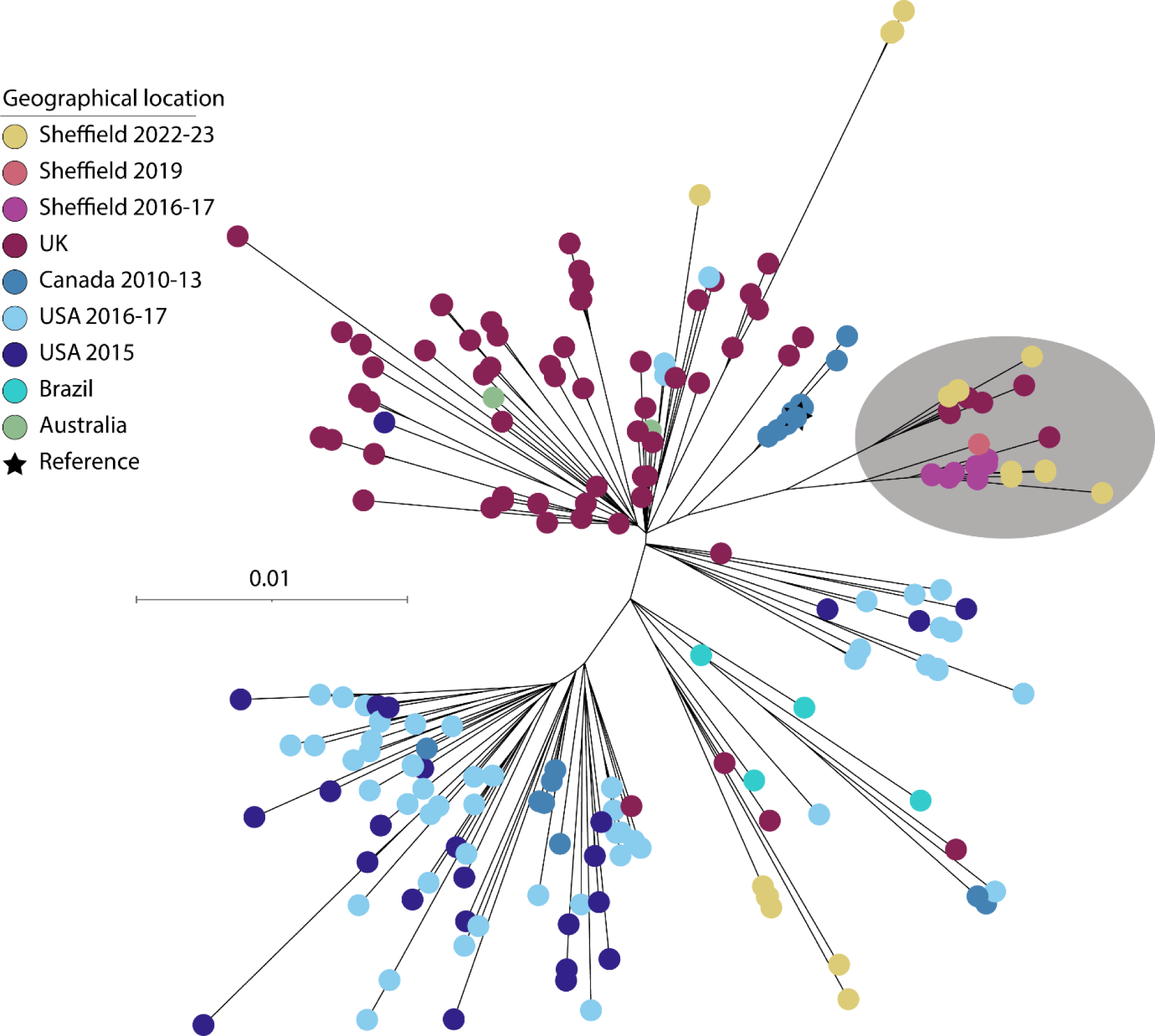
Phylogenetic analysis of Sheffield *emm*87 genomes collected in 2022-23 and 2016-17, within the context of global *emm*87 isolates. Alongside our Sheffield *emm*87 genomes from 2022-23 and 2016-17 we included publicly available *emm*87 genome data from the UK, 2001-18 (n=91) [31,38,39]; USA, 2015 (n=26) [45]; USA, 2016-17 (n=40) [46]; Canada, 2010-13 (n=22) [63]; Brazil, 2000-13 (n=4) [57]; and New Zealand, 2009-10 (n=2) [57]. A maximum likelihood phylogenetic tree was generated with the core gene alignment to reference strain NGAS743 (star). All isolates have a mutation in *hasA* that would truncate HasA but isolates in the lineage shaded in grey have an additional mutation in *hasB* that would truncate HasB. The scale bar represents the number of nucleotide substitutions per site. Two highly divergent UK strains were excluded from the tree for presentation purposes.

